# Water Network in the Binding Pocket of Fluorinated BPTI-Trypsin Complexes - Insights from Simulation and Experiment

**DOI:** 10.1101/2022.06.17.496563

**Authors:** Leon Wehrhan, Jakob Leppkes, Nicole Dimos, Bernhard Loll, Beate Koksch, Bettina G. Keller

**Affiliations:** Department of Biology, Chemistry, and Pharmacy, Freie Universität Berlin, Institute of Chemistry and Biochemistry, Arnimallee 22, Berlin, 14195 Germany; Department of Biology, Chemistry, and Pharmacy, Freie Universität Berlin, Institute of Chemistry and Biochemistry, Arnimallee 20, Berlin, 14195 Germany; Department of Biology, Chemistry, and Pharmacy, Freie Universität Berlin, Institute of Chemistry and Biochemistry, Takustr. 6, Berlin, 14195 Germany

## Abstract

Structural waters in the S1 binding pocket of *β*-trypsin are critical for the stabilization of the complex of *β*-trypsin with its inhibitor bovine pancreatic trypsin inhibitor (BPTI). The inhibitor strength of BPTI can be modulated by replacing the critical lysine residue at the P1 position by non-natural amino acids. We study BPTI variants in which the critical Lys15 in BPTI has been replaced by *α*-aminobutyric acid (Abu) and its fluorinated derivatives monofluoroethylglycine (MfeGly), difluoroethylglycine (DfeGly) and trifluoroethylglycine (TfeGly). We investigate the hypothesis that additional water molecules in the binding pocket can form specific non-covalent interactions to the fluorinated side chains and thereby act as an extension of the inhibitors. We report potentials of mean force (PMF) of the unbinding process for all four complexes and enzyme activity inhibition assays. Additionally, we report the protein crystal structure of the Lys15MfeGly-BPTI-*β*-trypsin complex (pdb: 7PH1). Both, experimental and computational data, show a step-wise increase in inhibitor strength with increasing fluorination of the Abu side chain. The PMF additionally shows a minimum for the encounter complex and an intermediate state just before the bound state. In the bound state, the computational analysis of the structure and dynamics of the water molecules in the S1 pocket shows a highly dynamic network of water molecules that does not indicate a rigidification or stabilizing trend in regards to energetic properties that could explain the increase in inhibitor strength. The analysis of the enthalpy and the entropy of the water molecules in the S1 binding pocket using Grid Inhomogeneous Solvation Theory confirms this result. Overall, fluorination systematically changes the binding affinity but the effect cannot be explained by a persistent water network in the binding pocket. Other effects, such as the hydrophobicity of fluorinated amino acids and the stability of the encounter complex as well as the additional minimum in the potential of mean force in the bound state, likely influence the affinity more directly.

**TOC GRAPHIC:** 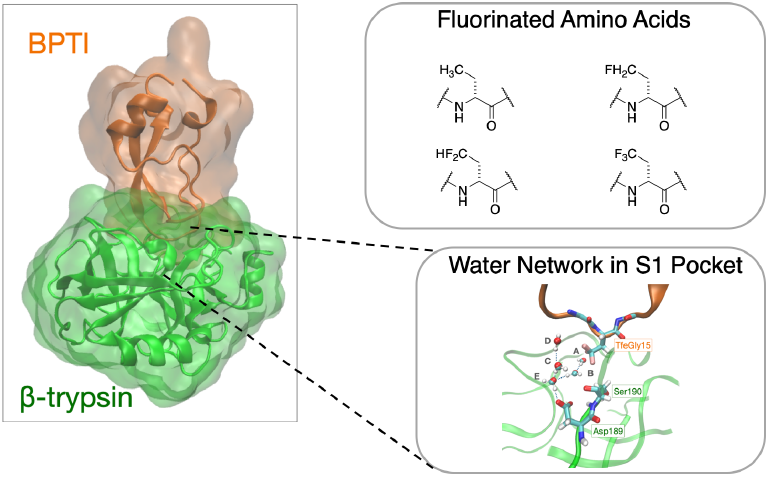

## I. INTRODUCTION

Hydrolysis of peptide bonds catalyzed by proteases is an ubiquitous process in all forms of life. This makes proteases important drug targets. However, drug design for these enzymes is difficult, because in proteases structural water molecules and hydration effects tend to play a critical role in ligand recognition and binding affinity^1–4^.

*β*-trypsin is a serine protease that acts as digestive enzyme in the small intestine of mammals where it breaks down food proteins. It is itself a drug target, but it also serves as model system for the study of protease inhibitors. In serine proteases, the enzymatic hydrolysis of the peptide bond in the substrate proceeds via a catatytic serine residue^5–8^, which is deprotonated by the combined action of an adjacent histidin and an aspartic acid. Deprotonation of the serine residue turns it into a strong nucleophile that can attack the carboxylic carbon atom in the peptide bond. The three critical residues serine, histidine and aspartic acid are called catalytic triad and are highly conserved in serine proteases^9^. In all serine proteases, the catalytic triad is located at the rim of a deep binding site called S1 binding pocket. Substrate specificity is realized by the amino acids within this binding pocket^9^. *β*-trypsin contains a negatively charged aspartic acid at the bottom of the S1 binding site. It thus recognizes positively charged amino acids, such as lysine and arginine^9^, and hydrolyses the peptide bond at their C-terminal side^9^.

*β*-trypsin is inhibited by multiple small molecules^10,11^, most notably benzamidine^12–14^. Additionally, several proteins have evolved as natural inhibitors of serine proteases in general, and *β*-trypsin in particular. Among them, Bovine pancreatic trypsin inhibitor (BPTI) is likely the most studied protein trypsin inhibitor^5–8,15–19^. BPTI also inhibits other serine proteases, and has been used under the drug name “aprotinin” to prevent excessive blood loss during surgery by acting on serine proteases involved in blood coagulation^20^.

BPTI is a small protein with only 58 residues and a molecular weight of 6.5 kDa^20^. It acts as a competitive inhibitor by inserting a lysine residue (P1 residue) into the S1 binding pocket and thus blocking the active site of *β*-trypsin. Interestingly, the lysine side-chain is too short to form a direct salt-bridge with the aspartic acid at the bottom of the S1 binding site. Instead, the binding is mediated by three water molecules that are trapped in the binding pocket^16,18^. One of the water molecules is located between the positively charged amino group of lysine and the negatively charged carboxyl group of aspartate. It accepts a hydrogen bond from the amino group and donates a hydrogen bond to the carboxyl group, and, via this hydrogen-bond network, bridges the gap between lysine in BPTI and aspartate in *β*-trypsin.

Despite the water-mediated binding mode, wild-type (wt) BPTI-*β*-trypsin complex is one of the strongest protein-protein complexes found in nature, with a binding constant^15,17^ of 5 to 6. 10^-14^ M and a corresponding binding free energy at *T* = 300 K of Δ*G*_bind_ = −76 kJ/mol. But BPTI is not very specific, and also binds to several other proteases^15,17^. By mutating the P1 residue in BPTI, i.e. the lysine that inserts into the binding pocket of *β*-trypsin, one can tune the binding free energy between −76 kJ/mol and −20 kJ/mol^21–23^. However, the BPTI variants remain promiscuous.

One way to affect the specificity of a protein inhibitor is by introducing non-natural amino acids^24^. In small molecule inhibitors the use of fluorine substituents has become a valuable tool to tune affinity and selectivity^25–28^. In Ref. 29, some of us introduced fluorinated non-natural amino acids at the P1 position in BPTI and investigated the effect on the inhibitor strength and the structure of the complex. Substituting lysine by the shorter and aliphatic residue *α*-aminobutyric acid (Abu) drastically reduced the inhibitor strength. This is expected, because the short side chain of Abu only reaches to about mid-way in the S1 binding pocket, and because it is an aliphatic side-chain, it cannot be stabilized by hydrogen bonds or charge interactions in the binding pocket. However, when two fluorine substituents were added to the terminal methyl group of Abu (Lys15DfeGly-BPTI), the inhibitor strength increased. It increased even further when the methyl group of Abu was fully fluorinated (Lys15TfeGly-BPTI).

The crystal structures of Lys15Abu-BPTI-*β*-trypsin complex showed five trapped water molecules in the S1 binding pocket. Three water molecules occupied the same positions as in the wt-BPTI-*β*-trypsin complex, the other two water molecules were found in the space that had been generated by truncating the P1 residue. The crystal structure of the two fluorinated complexes also showed five water molecules at the same positions, however with much lower B-factors, which indicates reduced mobility. This suggested the following mechanism for the increased affinity of the fluorinated BPTI-variants: The fluorine substituents form specific non-covalent interactions with the two additional water molecules, which are not possible with purely aliphatic side-chains. The two water molecules can thus act as an extension of the inhibitor and mediate binding to *β*-trypsin. In short: the water-mediated binding in the wt-BPTI-*β*-trypsin complex might be extended due to the presence of fluorine substituents^29^.

Crystal structures only provide static pictures of the complexes. On the other hand, molecular dynamics (MD) simulations can sample the structural fluctuations of the proteins as well as the dynamics of the water molecules in the binding pocket. Here, we use a combination of MD simulations, inhibition assays, and X-ray crystallography to elucidate the interaction of the fluorinated BPTI with trypsin and the water molecules in the binding pocket.

## II. METHODS

### A. Parametrization of the fluorinated Abu-variants

#### a. Parametrization protocol

The AMBER14sb^30^ force field was amended to include the parameters for the non-standard fluorinated amino acids and Abu. The parametrization process followed the protocol of Robalo et al. ^31^. Bonded parameters were generated from GAFF with the Acpype^32^ software. Lennard-Jones parameters for fluorine and hydrogen of fluoromethyl groups (H_*F*_) were taken over from Robalo et al.^31,33^. Atomic partial charges were calculated using a two stage RESP fitting protocol. For each amino acid, two dipeptide starting structures Ace-[Abu-variant]-Nme were constructed. One starting structure with the *φ* and *ψ* backbone dihedrals corresponding to alpha helix and the other one corresponding to beta sheet secondary structure. The structures were geometrically optimized using a molecular mechanics steepest descend algorithm. A single point QM energy calculation was performed at the HF/6-31-G* level of theory and the electrostatic potential (ESP) surface was evaluated using Gaussian16^34^. Using the Antechamber^35^ package, the ESP was fitted in a two stage RESP^36^ procedure to obtain atomic partial charges. During the RESP fit, atomic charges of the backbone amine and carbonyl were fixed to the AMBER14sb charge of the corresponding atoms for amino acids with neutral sidechains. The atomic partial charges of the Ace- and Nme-caps were fixed to the partial charges in Cornell et al.^37^. MD simulations of 100 ns each with constrained backbone atom positions were used to generate 200 conformations of each amino acid (100 conformations of alpha helix and 100 conformations of beta sheet). These conformations were submitted to a second RESP fit iteration under the same conditions as the first fit. The final atomic partial charges were calculated as the mean values of the second RESP fits.

#### b. MD simulations for second RESP fit iteration

The starting structures were placed into a cubic box of tip3p^38^ water with a buffer length of 1.0 nm between the box edges and the solute. The simulations were conducted in the modified AMBER14sb force field with atomic partial charges from the first iteration. The system was energy minimized using a steepest descent algorithm and equilibrated in the NVT ensemble for 100 ps at 300 K, followed by an equilibration in the NPT ensemble for 100 ps at 300 K and at 1.0 bar. Production MD simulations of the equilibrated system were performed in the NPT ensemble for a length of 100 ns. A velocity-rescaling scheme^39^ was used as thermostat and the Parinello-Rahman barostat^40^ was applied. The backbone atoms were constrained throughout equilibration and production simulation to conserve the backbone dihedral structure. Out of the resulting trajectories, 100 structures of the alpha helix conformations and 100 structures of the beta sheet conformations were extracted in equally spaced time intervals of 1 ns.

### B. MD simulations of the BPTI-Trypsin complexes and analysis

#### a. Unbiased simulations

MD simulations of the protein complexes were run with Gromacs 19-4^41–44^ and the self-parametrized amended AMBER14sb force field described above. The crystal structures of the protein complexes (pdb code: 4Y11, 4Y10, 4Y0Z) were freed from water, co-solutes and ions and placed into a dodecahedric box with 1.0 nm buffer between the solute and the box edges. One of the fluorine atoms in 4Y10 was mutated to a hydrogen atom to yield the Lys15MfeGly-BPTI *β*-trypsin complex. The complexes were solvated in tip4p^38^ water, energy minimized with a steepest descent algorithm and equilibrated in the NVT ensemble for 100 ps at 298 K, followed by an equilibration in the NPT ensemble for 100 ps at 298 K and 1.0 bar. A velocity-rescaling scheme^39^ was applied as thermostat and the Parrinello-Rahman barostat^40^ was applied. An initial simulation of 100 ns was run to extract starting conformations for the production simulations in equally spaced timesteps of 10 ns to ensure decorrelation. The extracted starting conformations were prepared, solvated, minimized and equilibrated as described above. 10 production simulations of 100 ns were run for each complex in the NPT ensemble at 298 K and 1.0 bar. Snapshots were saved in 1 ps timesteps. After production run, the trajectories were aligned based on backbone RMSD with the pdb structure 4Y11 as reference. Further handling of the trajectories for analysis was generally done with the MDTraj^45^ package.

To test the effect of the protonation state of His57 on the water dynamics, we produced one set of simulations with N(*δ*) and N(*ϵ*) of His57 protonated and one set of simulations where N(*δ*) is unprotonated.

#### b. Umbrella Sampling

*Fig.1b* Umbrella Sampling simulations of the complexes of the four Abu-derived BPTI-*β*-trypsin complexes were run in 39 windows along the reaction coordinate, each with a production simulation time of 50 ns. The reaction coordinate 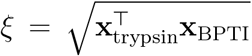 is the Euclidean distance between the heavy atom center of mass of *β*-trypsin, x_trypsin_, and the heavy atom center of mass of the BPTI-variants, x_BPTI_. Starting structures for the umbrella sampling windows were extracted from pulling simulations where a harmonic potential with a spring constant of *k*_pull_ = 1000 kJ · mol^-1^ · nm^-2^ was attached to the BPTI derivative and moved away from *β*-trypsin (increasing the center of mass distance) at a speed of 0.01 nm/ps. For the pulling simulations, the same pdb starting structures as in the unbiased MD runs of the complexes were freed from water, co-solutes and ions and placed into cubic box with 2.0 nm buffer between the solute and the box edges to ensure enough space for the pulling. The systems were energy minimized with a steepest descend algorithm and equilibrated in the NVT ensemble for 100 ps at 298 K, followed by an equilibration in the NPT ensemble for 100 ps at 298 K and 1.0 bar using the same thermostat and barostat as in the unbiased simulations. The starting structures for the umbrella sampling windows were extracted from the pulling simulations every 0. 05 nm along the reaction coordinate starting at 2.6 nm. These starting structures were equilibrated before the production runs of the umbrella sampling first in the NPT ensemble for 100 ps with the Berendsen barostat, then followed by a 1 ns equilibration using the Parinello-Rahman barostat. A harmonic potential with a force constant *k* = 6278 kJ · mol^-1^ · nm^-2^ was applied to the sampling window simulations and the production runs were conducted for the simulation time of 50 ns. Potential of mean force (PMF) profiles were calculated following the weighted histogram analysis method^46^ (WHAM) with the Gromacs-internal WHAM program. To estimate statistical uncertainty, a simplified bootstrapping scheme was applied by separating the trajectory of every window into five parts of 10 ns and combining the parts in the following way (beginning and starting time indicated in ns): (0-10, 10-20, 20-30, 30-40); (0-10, 10-20, 20-30, 40-50); (0-10, 10-20, 30-40, 40-50); (0-10, 20-30, 30-40, 40-50); (10-20, 20-30, 30-40, 40-50). PMF profiles of the bootstrapping samples were calculated and the mean and standard deviation were calculated. See supplementary material.

**FIG. 1.**
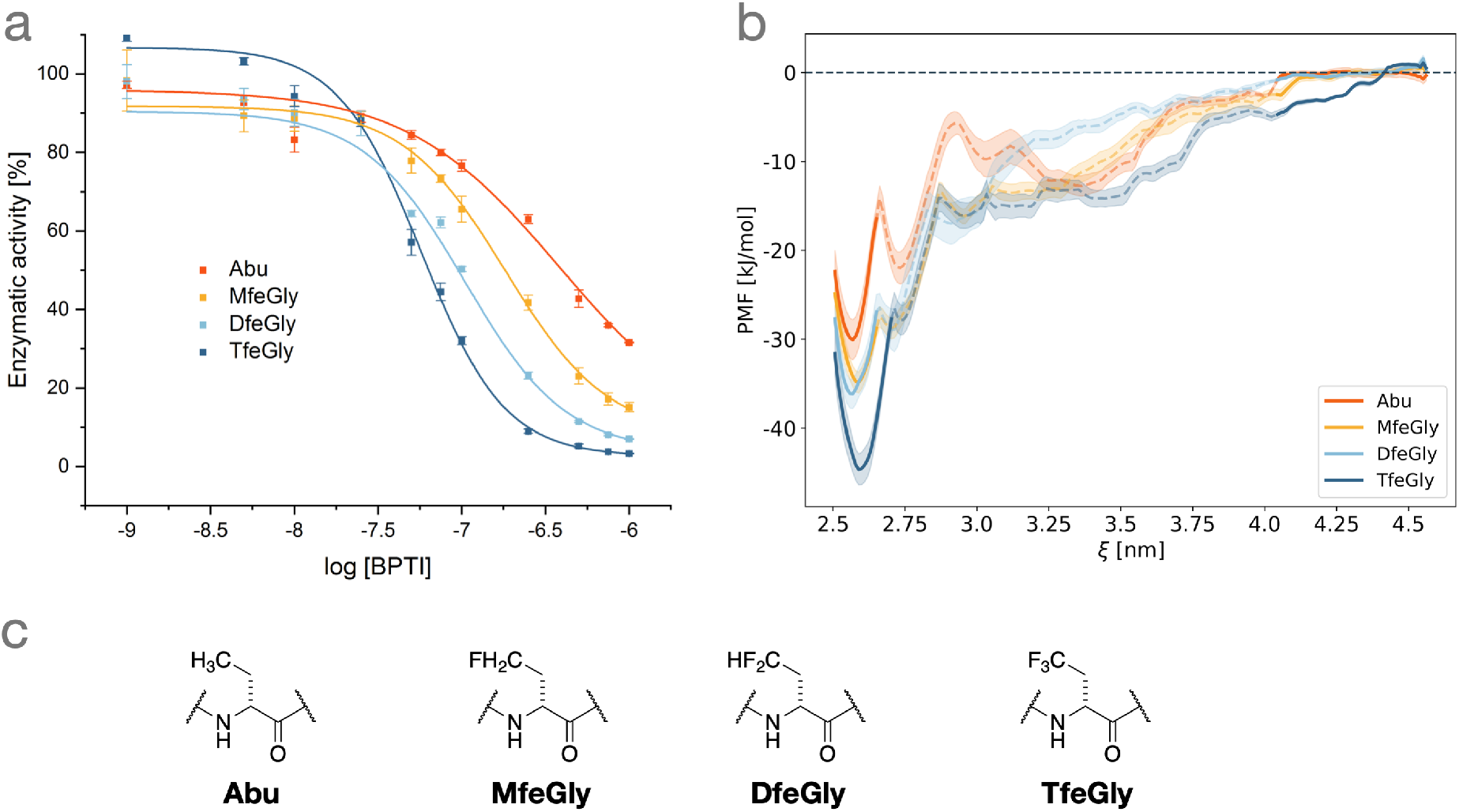
Inhibition of *β*-trypsin by BPTI variants with Abu derivatives at P1 position. (a) The inhibition assay shows a stepwise increase of inhibitor strength with increasing fluorination of the Abu side chain. (b) The Potential of Mean Force, here as an estimation of the binding free energy, from umbrella sampling simulations of the unbinding process shows the same trend. We here compare the depth of the free energy well of the fully bound state with the free energy of the unbound state. The unbinding path is shown as a dashed line as it shows one possible path of unbinding and should not be interpreted as a comprehensively sampled free energy profile of this region. (c) Chemical structures of Abu, MfeGly, DfeGly and TfeGly.

#### c. RMSF of long resident waters

*Fig. 3* Using the assignment of water molecules to the hydration sites A-E, water molecules that reside longer than 200 ps at a hydration site were identified. The RMSF of these water molecules, while they reside at the respective site was calculated and the average was estimated for every complex and hydration site.

**FIG. 2.**
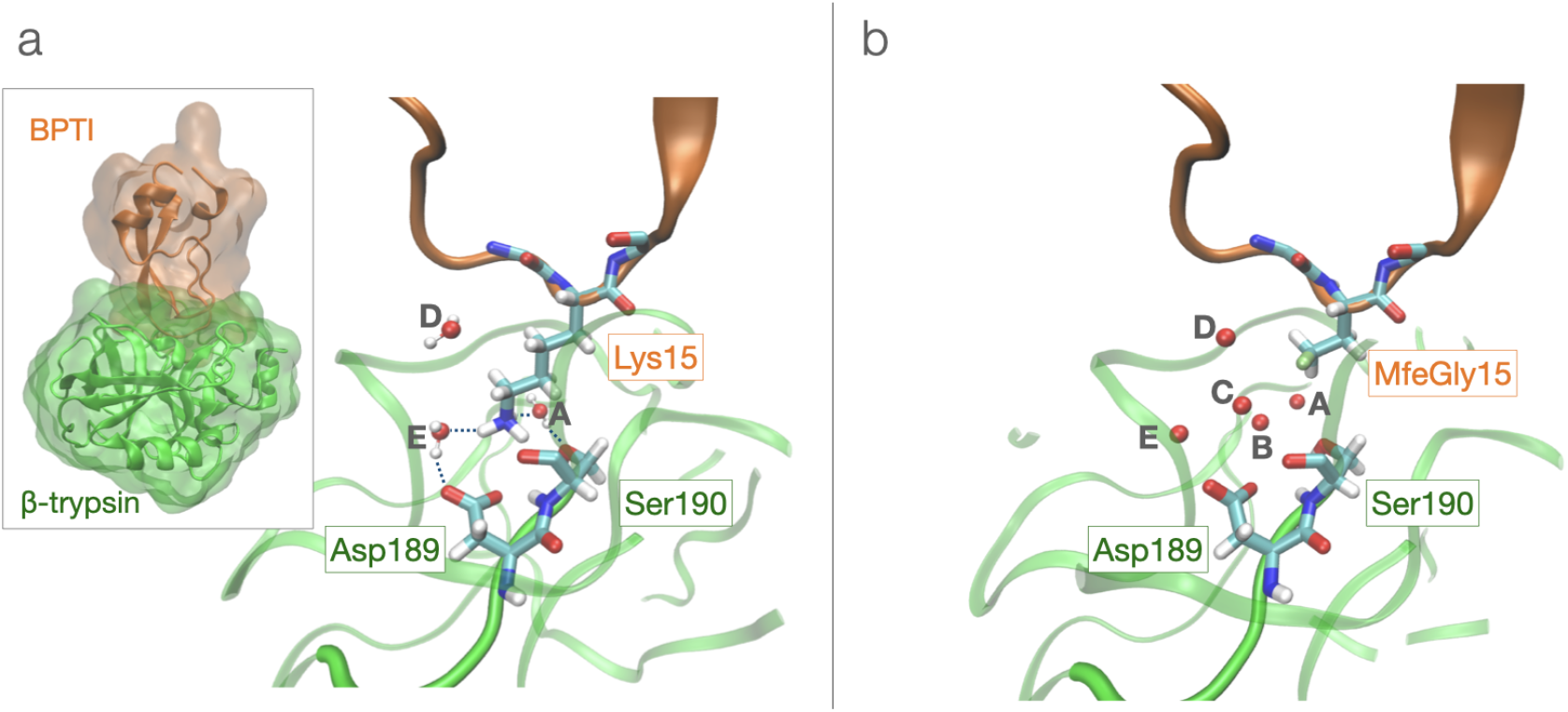
Structure of the BPTI-*β*-trypsin complex. (a) Wild-type BPTI-*β*-trypsin complex (pdb: 3OTJ^18^). (b) Lys15MfeGly-BPTI-*β*-trypsin complex (this work, pdb: 7PH1).

**FIG. 3.**
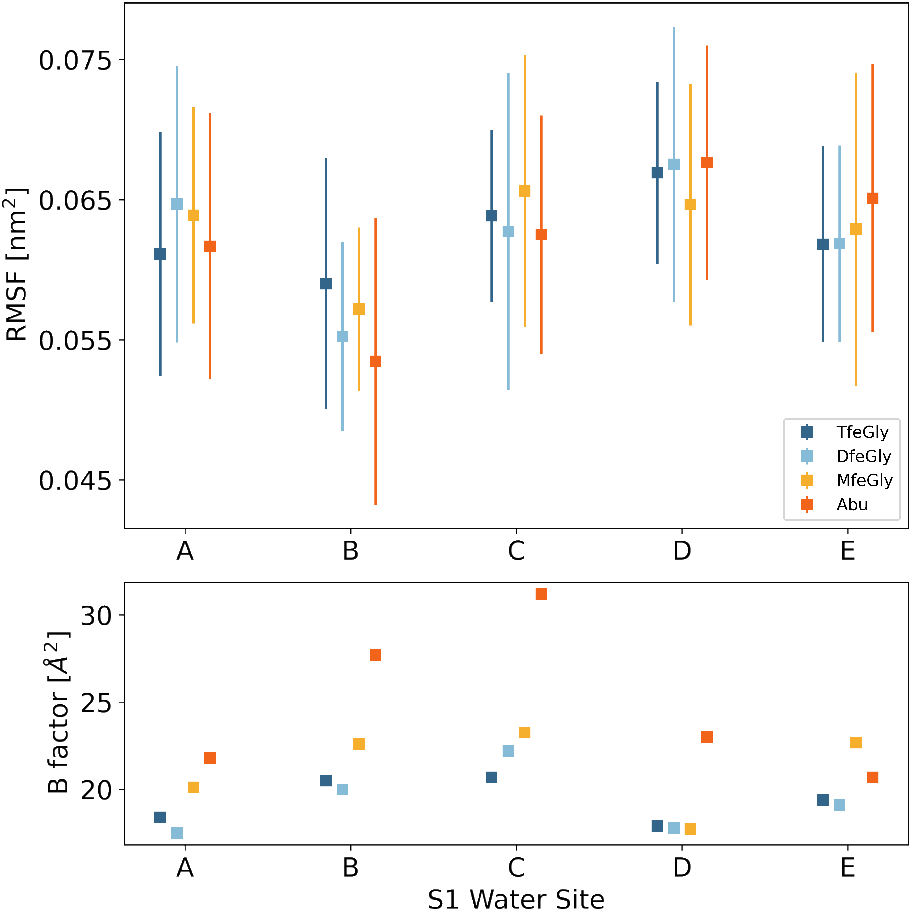
(top) RMSF of water molecules with a lifetime longer than 200 ps at the hydration sites A-E in the *β*-trypsin S1 binding pocket. The error bars indicate the standard deviation of the RMSF. (bottom) B factors of the waters in the *β*-trypsin S1 binding pocket as reported in ref. 29.

#### d. Assignment of water molecules to hydration sites

*Fig. 4b* As the water molecules which are present at the S1 pocket hydration sites A-E exchange frequently throughout the MD simulations, one specific water molecule was assigned to each site on a frame-by-frame basis. The position of the hydration sites was based on the positions of the water oxygens in the respective pdb structure of the protein complex. A sphere around each of the positions of these water oxygens at site A-E with a radius of 2.0 Å was defined and kept fixated at this position throughout the trajectory. Iterating over the trajectory frames, the water molecules whose oxygen atom is placed inside the spheres were identified. As the spheres for A-E were overlapping, in case a water molecule appeared in more than one sphere, it was assigned to the one where it was closest to the center (Voronoi tesselation). The closest water to the center within each sphere was assigned to be present at the respective hydration site for the frame if there were more than one water inside the sphere. If there was no water in the sphere after Voronoi tesselation, the hydration site was regarded as unoccupied. The code for the assignment of water molecules is available at: https://github.com/bkellerlab/track_waters.

**FIG. 4.**
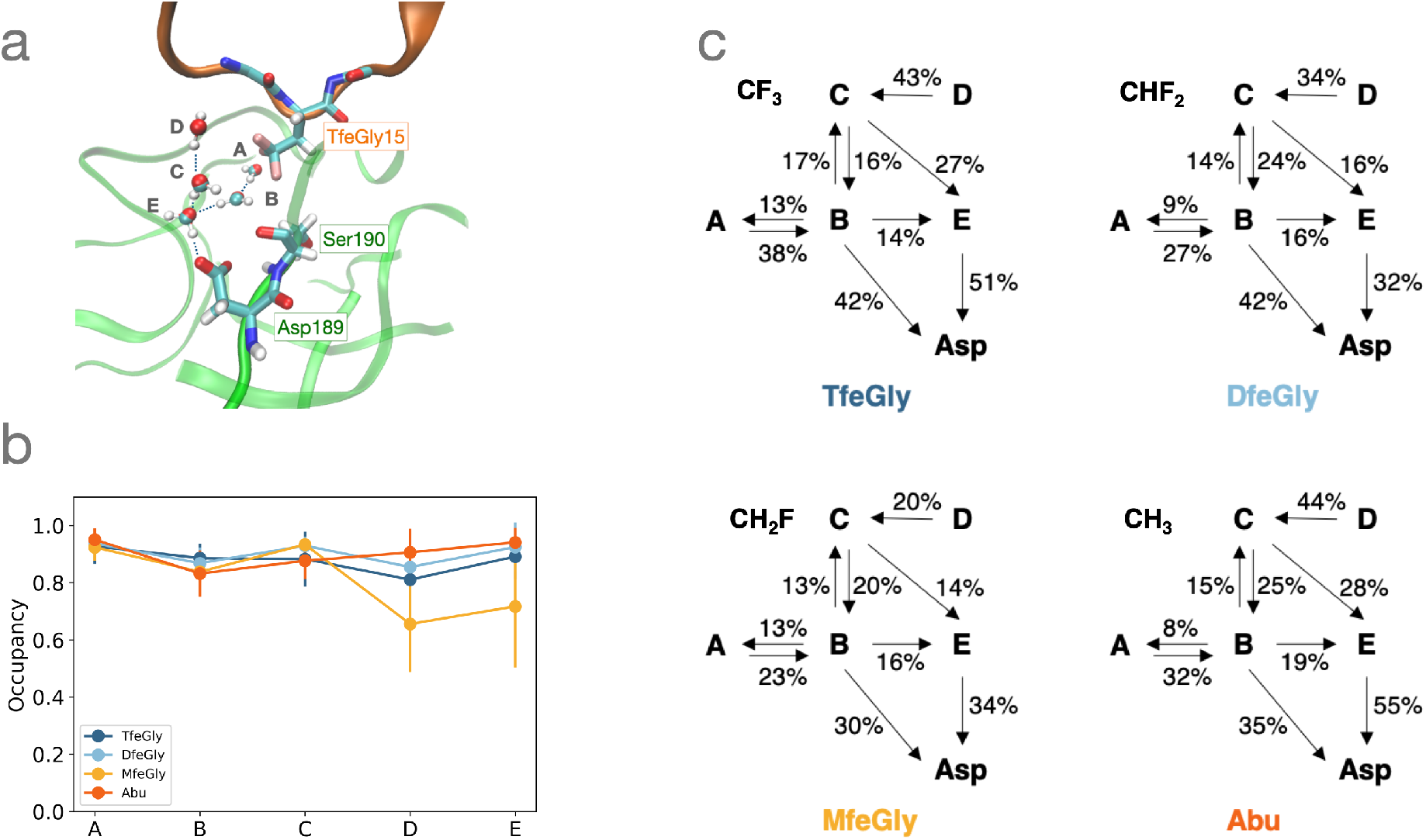
Structure of the water molecules in the S1 binding pocket. (a) Simulation snapshot of the S1 binding pocket with waters A-E. (b) Occupancy of the hydration sites A-E, showing how often the respective hydration site is populated by at least one water molecule throughout the simulations. (c) Hydrogen bond network. Relative populations of the hydrogen bonds formed by waters A-E to each other and to the side chain of Asp189. The arrows point from the donor to the acceptor.

#### e. Hydrogen bond network

*Fig 4c* To minimize the need for large amounts of memory, water that was not in proximity to the S1 binding pocket was deleted from the trajectories before submitting them to the calculation of hydrogen bonds. The water was removed dynamically on a frame-by-frame basis. The code is available at: https://github.com/bkellerlab/track_waters. Hydrogen bonds were estimated with the Wernet-Nilsson^47^ method of the MDTraj^45^ package. For every trajectory frame in particular, it was estimated if there are hydrogen bonds found between the waters at the hydration sites and to Asp189. The amount of frames in which the hydrogen bond was present was divided by the total number of trajectory frames to estimate the percentage how frequent the respective hydrogen bond is in place. To find hydrogen bond like interactions between the fluorinated methyl groups of MfeGly, DfeGly and TfeGly and the S1 waters A-E, the distance between their oxygen atoms and the fluorine atoms, as well as the O-H-F angles were calculated for every trajectory frame. Using the definition in ref. ^48^ of a hydrogen bond like interaction, the frames in which the distance was smaller than 0.35 nm and the corresponding angle larger than 160° were counted as showing a hydrogen bond like interaction between fluorine and the respective water molecule. In a separate analysis, the distance and O-H-F angle of every water molecule coming close enough to the fluorine atoms was calculated for the trajectories to find the number of frames in which hydrogen bond like interactions are in place involving any water molecule, not just those that could be assigned to the S1 pocket hydration sites.

#### f. Survival probability in the S1 pocket

*Fig 5a* The survival probability of water molecules in the S1 binding pocket was calculated using the waterdynamics^49^ module of the MDAnalysis^50,51^ package. The volume that belongs to the S1 pocket was defined as 5Å-spheres around the side chain ends of Asp189 and Ser190. The survival probability of a water molecule at time *t* in the S1 pocket, given for a lag-time *τ*, is defined as the probability that it is still present in the pocket at time t + *τ*. The probability is averaged over the trajectory:

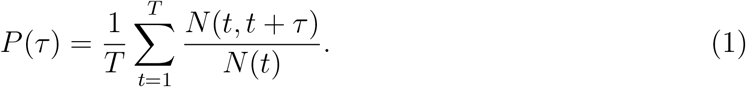

**FIG. 5.**
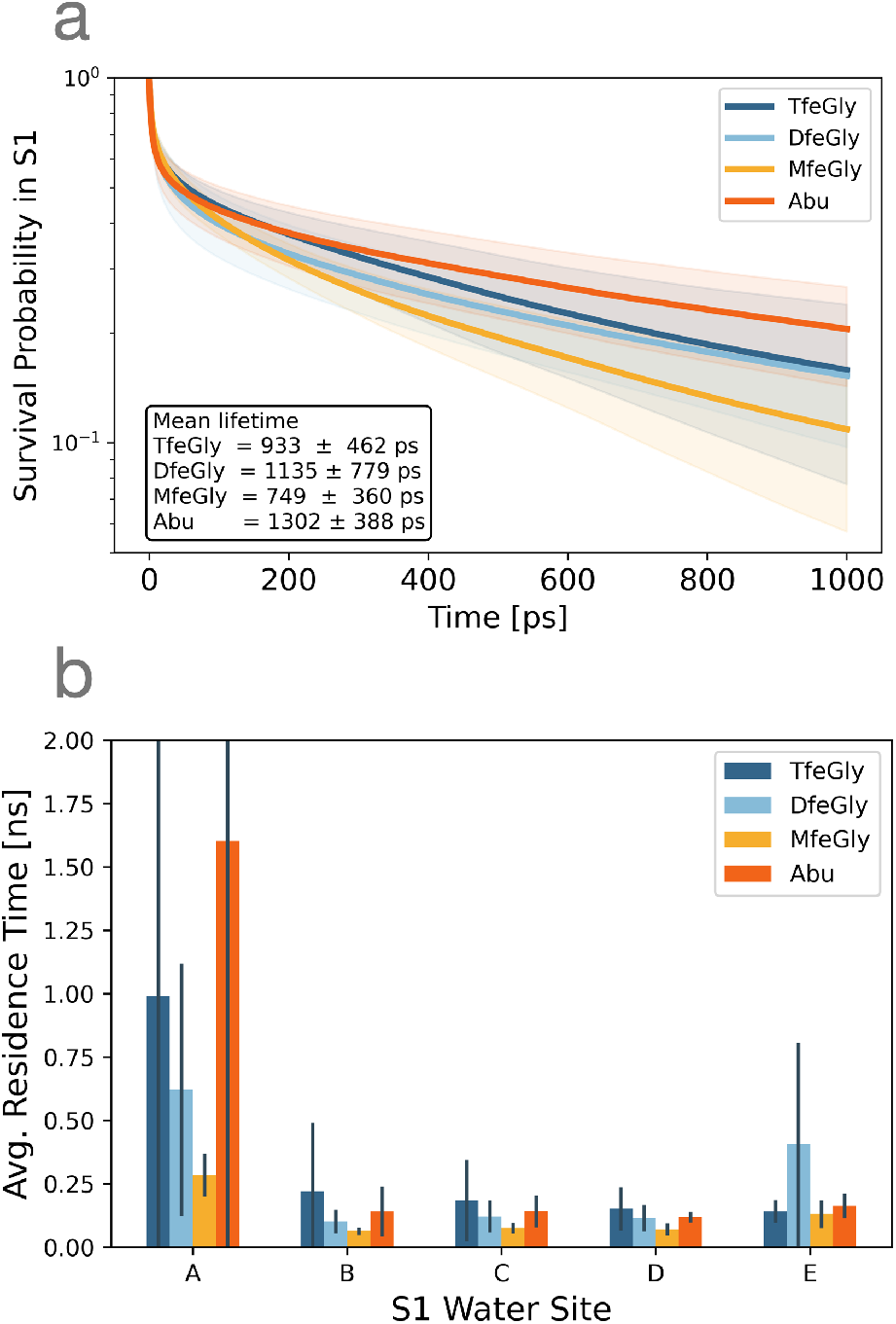
Dynamics of the water molecules in the S1 binding pocket of the *β*-trypsin BPTI complexes. (a) Average survival probability of an arbitrary water molecule in the S1 binding pocket. The mean lifetime is approximated by fitting a single-exponential decay function to the data after the initial burn phase. (b) Average residence time of a water molecule at the hydration sites A-E. The data is calculated from a total of 1 μs of simulation data for each of the protein complexes.

*T* is the total number of trajectory frames, *N*(*t*) is the number of water molecules in the region of interest at the time and *N*(*t*, *t* + *τ*) is the number of water molecules that remain in the region of interest after the lag-time *τ*. The survival probability up to a lag-time of 1 ns was calculated for each trajectory. An average was formed from the results for each complex and plotted in Fig. 5a. The mean lifetime t was approximated by fitting a single-exponential decay function of the form 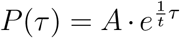 to the survival probability of each trajectory after an initial burn phase of 100 ps to account for waters that have very short survival lifetime due to fluctuating in and out around the edge of the defined S1 pocket zone.

#### g. Mean residence time

*Fig 5b* After assigning water molecules to the S1 pocket hydration sites as described above, exchanges of water molecules were counted. The total simulation time was divided by the number of exchanges to estimate the mean residence times of a water molecule at the particular hydration sites. To account for fluctuations on the edge of the spheres, simulation chunks of 20 ps were regarded, where the water molecule found most often at the hydration site in the chunk was assigned to that site for the whole chunk. The resulting mean residence times were averaged over the trajectories for each complex.

#### h. GIST calculation

*Fig. 6* Thermodynamic properties of water molecules were calculated based on Grid Inhomogeneous Solvation Theory (GIST). For the underlying simulations, the crystal structures of the protein complexes (PDB code: 4Y11, 4Y10, 4Y0Z) were prepared and solvated in the same way as the unbiased simulations of the BPTI-Trypsin complexes. The systems were equilibrated in the NVT ensemble at 300 K for 10 ns. Positional restraints were imposed on every protein heavy atom for the entire equilibration and production runs of 100 ns length in the NVT ensemble. The GIST calculations were performed with the SSTMap^52–55^ command-line tool. A cubic grid with side-length of 26 Å and grid spacing of 0.5 Å was centered on the C_*γ*_ atom of the BPTI P1 residue to capture the active site. The resulting voxel properties of E_*sw*_, E_*ww*_, dS_*tr*_T and dS_*or*_T were visually inspected in PyMol and clusters were identified around the S1 water hydration sites A-E of the crystal structures. Spheres of 2.0 A radius were defined around the positions of the hydration sites (defined by the position in the crystal structure of 4Y11) and the voxels inside the spheres were grouped to the corresponding water site. Due to overlapping of the spheres, voxels were assigned to the closest water sites by Voronoi tessellation. The thermodynamic properties 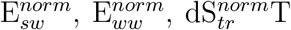 and 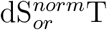 of the voxels were averaged by Boltzmann weighted averaging respectively for each voxel cluster group to obtain the mean magnitude of the thermodynamic property for a voxel assigned to the water site. The Boltzmann weights were determined by the number of water molecules *N*_wat_ from the SSTMap output. For each complex, four different simulations were equilibrated and run and the analysis was done on all four simulations to estimate statistical uncertainty.

**FIG. 6.**
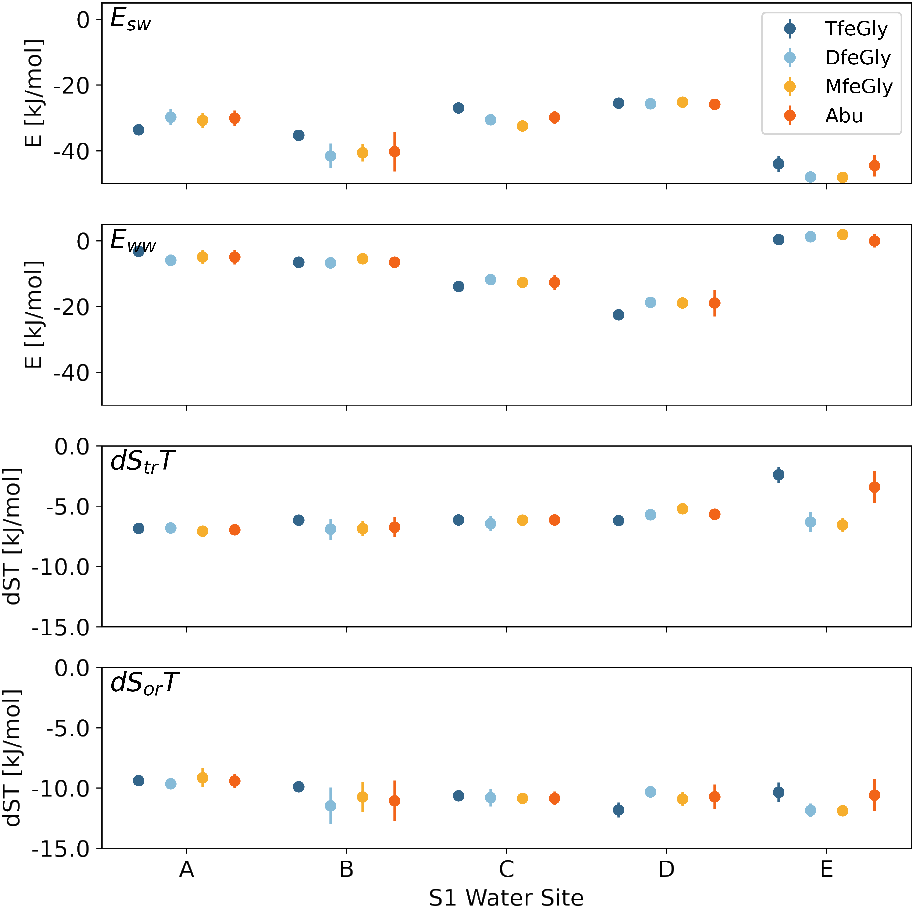
The solute-water (*E_sw_*) and the water-water (*E_ww_*) interaction energy and translational (*dS_tr_T*) and rotational (*dS_or_T*) entropy for S1 binding pocket waters, calculated from GIST.

#### i. Dihedral angle distribution

*Fig 7* The dihedrals *χ*_1_ and *χ*_2_ were calculated from the unbiased MD simulations with the MDTraj^45^ package. The histograms present an average distribution of the 10 trajectories for each complex. The standard deviation was added and substracted from the average to calculate the error lines. The dihedrals in the PDB crystal structures 4Y11, 4Y10, 7PH1 and 4Y0Z were measured for comparison to the dihedral distributions from the MD simulations.

**FIG. 7.**
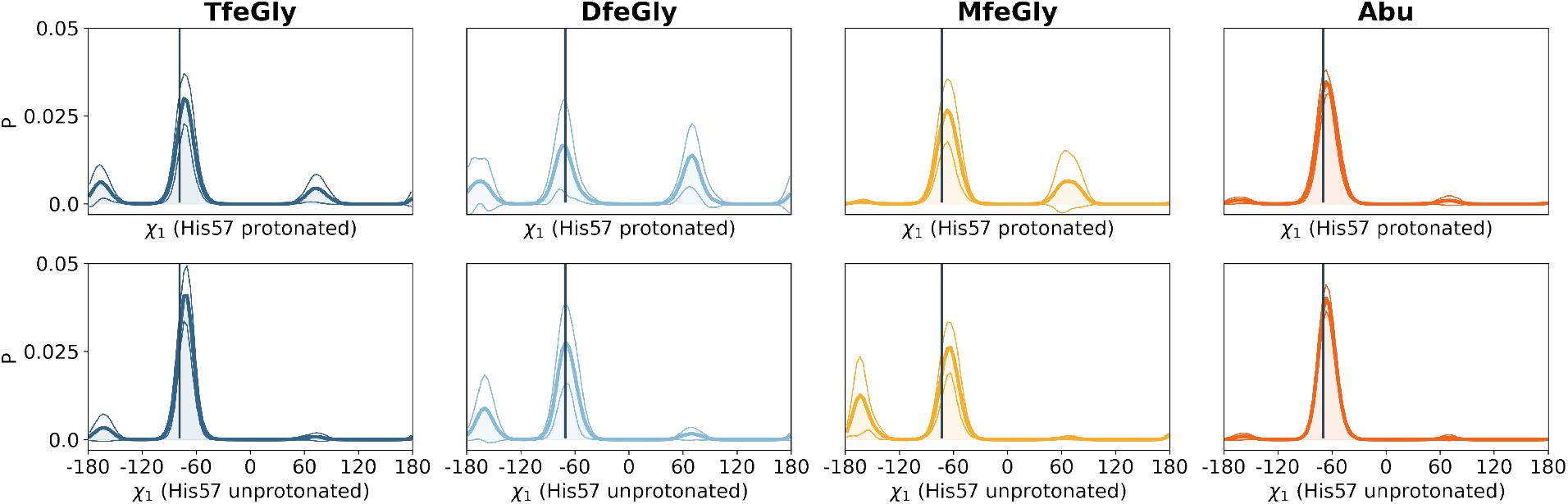
Influence of the protonation state of His57 on the *χ*_1_-torsion angles of the P1 residue. Histograms with the probability density P of the bins. The vertical line marks the value of the torsion angle from the respective pdb structure. *Top row:* His57 protonated. *Bottom row:* His57 unprotonated. Standard deviations are indicated as thin lines.

### C. Enzyme activity inhibition assay: Fig.1a and Tab. I

**TABLE I.** Binding constants and binding free energies derived from the inhibition assay. We assume *T* = 300 K.

| system | Ref. 29 |  | experiment, this work |  |  |
| --- | --- | --- | --- | --- | --- |
| | $K_a$ [$M^{-1}$] | $\Delta G_{\text{bind}}$ [kJ/mol] | $IC_{50}$ [M] | $K_a$ [$M^{-1}$] | $\Delta G_{\text{bind}}$ [kJ/mol] |
| Lys15Abu | | | $(3.83^{+0.102}_{-0.102}) \cdot 10^{-7}$ | $(2.91^{+0.3}_{-0.2}) \cdot 10^6$ | $-37.1^{+0.1}_{-0.2}$ |
| Lys15MfeGly | | | $(1.83^{+0.109}_{-0.109}) \cdot 10^{-7}$ | $(6.09^{+0.8}_{-0.5}) \cdot 10^6$ | $-39.0^{+0.2}_{-0.3}$ |
| Lys15DfeGly | $3.88 \cdot 10^7$ | -43.6 | $(1.05^{+0.115}_{-0.115}) \cdot 10^{-7}$ | $(1.06^{+0.2}_{-0.1}) \cdot 10^7$ | $-40.4^{+0.3}_{-0.5}$ |
| Lys15TfeGly | $5.20 \cdot 10^7$ | -44.3 | $(5.73^{+0.115}_{-0.115}) \cdot 10^{-8}$ | $(1.94^{+0.2}_{-0.1}) \cdot 10^7$ | $-41.9^{+0.2}_{-0.3}$ |

#### a. Enzyme activity inhibition assay

For inhibitory assays increasing concentrations of BPTI variants (0, 1, 5, 10, 25, 50, 75, 100, 250, 500 750, 1000 nm) were incubated with 20 μL *β*-trypsin (100nM, Sigma-Aldrich) in 200mM Triethanolamine, 20mM CaCl_2_, pH 7.8 in a 96-well-plate for 1 h. 20 μL N*α*-Benzoyl-L-arginine 4-nitroanilide hydrochloride (BApNA, Sigma-Aldrich) (1mM) were added to 180 μL preincubated enzyme/inhibitor mixture. Hydrolysis of BApNA and formation of the product p-Nitroaniline (pNA) was monitored by measuring absorbance at 405 nm in an Infinite M200 microplate reader (Tecnan Group AG, Männedorf, Switzerland) for 30 min at 25 °C. Initial velocities were determined by plotting absorbance against reaction time and IC_50_ values by plotting residual enzyme activity against logarithmic inhibitor concentration using OriginPro 2020b (OriginLab Corporation, Northampton, MA, USA).

#### b. Conversion between IC_50_ and ΔG_bind_

To relate the half maximal inhibitory concentrations (IC_50_) measured in the inhibition assays to the binding free energy Δ*G*_bind_, we consider the binding equilibrium of of the BPTI-variants Lys15X-BPTI to *β*-trypsin

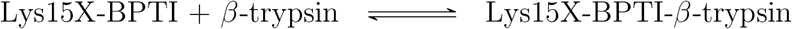

The enzyme inhibition constant, or inhibitor dissociation constant, *K_i_* for this equilibrium is defined to be equal to the dissociation constant of the inhibitor-enzyme complex

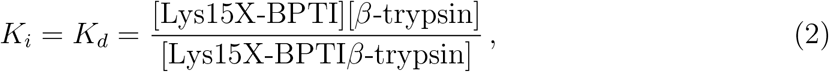

where [*A*] denotes the concentration of species *A*. Thus, *K_i_* has units of concentration. We related the half maximal inhibitory concentrations (IC_50_) measured in the inhibition assays to the inhibitor dissociation constant via the Cheng-Prusoff equation^56^

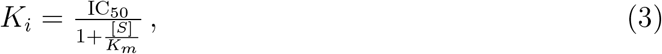

where *K_m_* = 8.85 ± 3.60 mM is the Michaelis constant of *β*-trypsin. We determined the Michaelis constant of *β*-trypsin ourselves (see below). The substrate concentration was [*S*] = 1 mM in the inhibition assay in Fig. 1. With these values IC_50_ and *K_i_* can be interconverted as *K_i_* = 0.898 · IC_50_ and IC_50_ = 1.113 · *K_i_*.

Table I reports the association constant which is the inverse of the inhibition constant

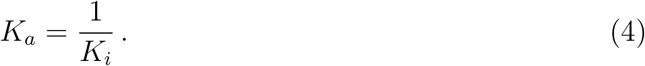

and the binding free energy

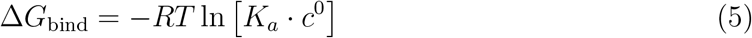

where *R* = 8.314 J/(K mol) is the ideal gas constant, *T* = 300 K is the temperature, and *c*^0^ = 1 M is the standard concentration. We obtained the uncertainties in *K_a_* and ΔG_bind_ by propagating the uncertainties in IC_50_ and *K_m_*.

#### c. Determination of Michaelis-Menten constant

To determine the Michaelis-Menten constant *K_m_*, 20 μL N*α*-Benzoyl-L-arginine 4-nitroanilide hydrochloride (BApNA, Sigma-Aldrich) were added in varying concentrations (0, 0.1, 0.2, 0.5, 0.7, 1.0, 1.2 mM; final concentrations of the measurement) to 180 μL *β*-trypsin (100 nM, final concentration, Sigma-Aldrich) in 200mM Triethanolamine, 20mM CaCl_2_, pH 7.8 in a 96-well plate. Hydrolysis of BApNA and formation of the product p-Nitroaniline (pNA) was monitored by measuring absorbance at 405 nm in an Infinite M200 microplate reader (Tecan Group AG, Männedorf, Switzerland) for 10 min at 25 °C. Initial velocities were determined by plotting absorbance against reaction time. *K_m_* was determined by using Enzyme Kinetics plug-in in OriginPro 2020b (OriginLab Corporation, Northampton, MA, USA).

### D. Protein Crystallography

To form complex between trypsin and BPTI variants, 60 μL of 2.0 mM BPTI in 25mM Tris, 10 mM CaCl_2_, pH 7.4 were added to 50 μL of bovine trypsin in 25 mM Tris/HCl, 10 mM CaCl_2_, pH 7.4. The solution was incubated for 12h at 4°C. *β*-trypsin-BPTI complex was purified by dialysis. Initial crystals were obtained by the sitting-drop vapor-diffusion method at 18 °C with a reservoir solution containing 2.2 M ammonium sulfate. Inter-grown crystals were used to prepare a seed stock. With a cat whisker, seeds were transferred to a freshly prepared crystallization drop. Well-formed crystals were soaked in 30 % (v/v) glycerol plus reservoir solution and frozen in liquid nitrogen. Data was collected at Berlin BESSYII, beamline 14.2. X-ray data collection was performed at 100 K. Diffraction data were processed with the XDS in space group I222 (Table SI-XX).^57^ The structure of the Trypsin-BPTI-MfeGly was solved by molecular replacement with the coordinates of Trypsin-BPTI as search model using PHASER^58^. The structure was refined by maximum-likelihood restrained refinement and TLS refinement^59^ using PHENIX^60,61^ followed by iterative, manual model building cycles with COOT^62^. Model quality was evaluated with MolProbity^63^. Figures were prepared using Visual Molecular Dynamics^64^. The atomic coordinates and structure factor amplitudes have been deposited in the Protein Data Bank under the accession code 7PH1 (Trypsin BPTI-MfeGly).

## III. RESULTS & DISCUSSION

### A. Parametrization of the fluorinated variants of aminobutyric acid (Abu)

The following fluorinated variants of aminobutyric acid (Abu) are included in this study: monofluoroethylglycine (MfeGly), difluoroethylglycine (DfeGly) and trifluoroethylglycine (TfeGly) (see Fig. 1.c). Fluorinated carbon atoms are not reproduced very accurately by the parameter sets of standard MD force fields. We therefore amended the AMBER14SB^30^ force field and customized the parameters for the-CFH_2_, −CF_2_H and −CF_3_ in MfeGly, DfeGly and TfeGly following the procedure suggested by Robalo et al.^31,33^ (see also section II). We generated the bonded parameters using the GAFF with the Acpype^32^ software. For the van-der-Waals interactions, we used the Lennard-Jones parameters for fluorine and hydrogen of fluoromethyl groups (H_*F*_) from Robalo et al.^31,33^, who had determined these parameters by fitting against hydration free energies and obtained very accurate results. For the Coulomb interactions, we calculated the atomic partial charges with a two stage RESP^36^ fitting protocol, similar as in Ref. 31. In the second stage of this protocol, a conformational sample is generated by an MD simulation with preliminary charges, and the charges are then optimized based on this sample. We used a large sample of 200 structures generated by 100 ns of MD simulations for each amino acid and backbone conformation to ensure that the conformational ensemble of each of the amino acids it well represented. The thus obtained partial charges are reported in SI Fig. 1 and agree well with the charges from Refs. 31 and 33. In our forcefield, the partial charge of fluorine is slightly less negative than in the references. As a consequence, the partial charge of the highly fluorinated C_*γ*_ of TfeGly and DfeGly is moderately less positive than in the references. The amended force field is included in the supplementary material.

### B. Inhibition assay and binding constants

Next, we confirmed that fluorination of Lys15Abu-BPTI increases the inhibitor strength by repeating the inhibition assay from Ref. 29. We now additionally had the monofluorinated variant Lys15MfeGly-BPTI available and included it in the assay (Fig. 1.a). Furthermore, we now could determine the IC_50_ value of the unfluorinated variant Lys15Abu-BPTI. As before, fluorination of the methyl group in the Abu side chain lead to a stepwise increase of the inhibitor strength from IC_50_ = 383 nM (3.83 · 10^-7^ M) for the unfluorinated Lys15Abu-BPTI to IC_50_ = 57 nM (5.7 · 10^-8^ M) for the fully fluorinated Lys15TfeGly-BPTI. The IC_50_ for Lys15DfeGly-BPTI and Lys15TfeGly-BPTI are in good agreement with the previously reported values^29^ (Tab. I).

We converted the IC_50_ values into binding constants and binding free energies (see Tab. I). This conversions depends on several assumptions, since the inhibition assay reports on enzyme activity and not on thermodynamic property^65,66^. However, within these assumptions, the binding free energy decreases almost linearly from −37.1 kJ/mol in Lys15Abu to −41.9 kJ/mol in the fully fluorinated Lys15TfeGly. Thus, the compounded effect of fluorinating the Abu sidechain is −4.8 kJ/mol, were each fluorine atom contributes about out −1.5 kJ/mol to stabilizing the fluorinated BPTI-*β*-trypsin complex. Overall, this is a small but reproducible and robust effect in the enzyme activity inhibition assay.

We find the same effect in our MD simulations. Fig. 1.b shows the potential of mean force (PMF) for the unbinding of the four complexes, which we calculated by umbrella sampling and weighted-histogram analysis (39 umbrella windows and 1.95 μs total simulation time per PMF curve, see section II). The reaction coordinate 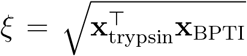 is the Euclidean distance between the center of mass of *β*-trypsin, X_trypsin_, and the center of mass of the BPTI-variants, x_BPTI_. The minima at ξ = 2.6 nm correspond to the fully bound state, and for ξ > 4.1 nm the complex is fully unbound. Assuming that the PMF of the unbound state of the four systems can be matched, we find that the PMF of the bound state decreases from the unfluorinated Lys15Abu-BPTI to the fully fluorinated Lys15TfeGly-BPTI, and thus reproduces the effect from the inhibition assay. Also, the relative depths of the PMF minima are similar in magnitude as the free-energy differences derived from inhibition assays.

We believe these trend in the PMFs to be robust, even though we do not report quantitative free-energy differences. In order to calculate absolute free energy differences from a PMF, the translational and rotational freedom of the unbound BPTI-variants needs to be taken into account accurately^67–69^. The translational and rotational freedom of the bound and unbound BPTI-variants are are analyzed and discussed in the supplementary material (Fig. S10-S13).

We used a single simulation of 50 ns per umbrella window. However, the sampled exit path and consequently the PMF are known to be sensitive to the starting points of the umbrella simulations. A well-converged transition region between bound and unbound state likely requires a sampling the PMF along additional reaction coordinates. Nevertheless we would like to point out several interesting features in Fig. 1.b. Most notably, the PMF of Lys15Abu-BPTI is more rugged than the PMFs of the fluorinated variants. Lys15Abu-BPTI has a broad minimum between ξ = 3.0 nm and the unbound state which corresponds to an encounter complex. There is a considerable barrier at ξ ≈ 3.0 between this encounter complex and the bound state. In the fluorinated variants, we instead find a slowly decreasing PMF between ξ = 3.0 nm and the unbound state, which is separated only by a minimal barrier from the bound state. Second, the fully bound state in Lys15Abu-BPTI (ξ = 2.6 nm) is separated by a sharp and large barrier from a pre-bound state at (ξ = 2.7 nm). The fluorinated variants also show such a pre-bound state. However, the pre-bound state is not as stable (i.e. higher PMF relative to the fully bound state) and only separated by a small barrier from the fully bound state.

### C. Crystal structure of the Lys15MfeGly-BPTI-*β*-trypsin complex

Fig. 2.a shows the structure of the wild-type(wt)-BPTI in complex with *β*-trypsin^18^. *β*-trypsin has a deep S1 binding pocket, and the catalytic triad is located at the rim of this binding pocket. See Ref. 16 for a schematic drawing of the binding pocket including the catalytic triad. Asp189 at the bottom of the binding pocket governs the specificity of *β*-trypsin for positively charged side chains, such as lysine and arginine. wt-BPTI inserts the residue Lys15 into the binding pocket (P1 residue). However, the lysine side chain is not long enough to form a direct salt bridge to Asp189. Instead, the contact is mediated by hydrogen bonds to a trapped water molecule (E). Lys15 forms additional hydrogen bonds to a second trapped water molecule (A) and to Ser190 in *β*-trypsin. Finally, crystal structures of BPTI-*β*-trypsin complexes show a third water molecule (D) in the binding pocket close to the backbone of Lys15. The water molecules in Fig. 2 are labeled according to Ref. 29.

Having synthetic access to the monofluorinated amino acid MfeGly^70^ and the corresponding variant of BPTI^71^, we now determined the crystal structure of the Lys15MfeGly-BPTI-*β*-trypsin complex and deposited it in the Protein Data Bank (pdb: 7PH1). The crystal structure shows five water molecules in the binding pocket (Fig. 2). Water molecules A, D and E are located a the same positions as in the wt-BPTI-*β*-trypsin complex. Because the side chains of the P1 residue in the Lys15MfeGly-BPTI variants is considerably shorter than lysine, there is more space in the S1 binding pocket. This additional space is occupied by two additional water molecules, labeled B and C. This is in line with the previoulsy published crystal structures of the Lys15Abu-BPTI-*β*-trypsin complex and its di- and tri-fluorinated variants^29^, which also show water molecules at the same five positions. See Ref. 29 for a detailed discussion of the distances between the trapped water molecules and the two proteins in the crystal structures.

As noted in Ref. 29, the B-factors of the water molecules in the Lys15Abu-BPTI-*β*-trypsin complex differ markedly from the B-factors in the fluorinated variants. The lower B-factors in the fluorinated variants indicate that the water molecules are less mobile in these variants and might act as a “hydration shell” or “contour line” of the fluorinated P1 residue which effectively acts as an extension of this residue^29^ and thereby increases the binding affinity. The B-factors of water molecules in the mono-fluorinated Lys15MfeGly-BPTI-*β*-trypsin complex complete this picture. They are slightly higher than the B-factors of the di- and tri-fluorinated variants, but considerably lower than the B-factors of the complex with Lys15Abu-BPTI (Fig. 3. (bottom)).

### D. Water network in the S1 binding pocket

#### a. The hypothesis

Based on the mobility of the water molecules in the crystal structures, the following mechanism to explain the influence of the fluorine substituents on the binding affinity seems plausible^29^: In contrast to Abu, the (partially) fluorinated methylgroup in MfeGly, DfeGly and TfeGly can form directed non-covalent interactions with water molecules B and C. These two water molecules form hydrogen bonds to water molecules E and A, and the resulting hydrogen bond network that bridges the gap between the truncated P1 side chain and Asp189. The stability of this hydrogen bond network increases with the number of fluorine atoms, and thereby increases the binding affinity.

With MD simulations, we can test this hypothesis by monitoring the structure, the dynamics and the energy of the water molecules in the S1 binding pocket. Specifically, we would expect to see the following trends with increasing fluorination:

- increasing stability of the hydrogen bond network in the binding pocket
- decreasing fluctuations in the hydrogen bond network in the S1 binding pocket.
- increasing survival times of the water molecules in the binding pocket
- decreasing mobility of the water molecules within the binding pocket
- decreasing entropy of the waters in the S1 binding pocket
- a systematic trend in the water-water and the water-solute enthalpy

#### b. Structure of the water molecules in the S1 binding pocket

In all four complexes, the five positions A-E are occupied on average between 84 % and 95 % of the simulation time. The only exception is the Lys15MfeGly-BPTI-*β*-trypsin complex, in which positions D and E are populated less frequently (Fig. 4.b). Thus, on average the S1 water network of the solvated complexes resembles the water network in the crystal structures.

The waters in the narrow S1 pocket are in direct proximity to each other and can easily form hydrogen bonds. We calculated the frequency of the hydrogen bonds between waters A-E by counting in how many snapshots of our simulations a hydrogen bond is accepted from or donated to another of the S1 waters or Asp189 of *β*-trypsin. Hydrogen bonds were defined using the Wernet-Nilsson^47^ method as implemented in the MDTraj^45^ package, which corresponds to identifying strong hydrogen bonds (compare Ref. 47 and table 1.5 on p.13 in Ref. 72). We find the same hydrogen bond network in all of the variants, with slightly to moderately varying relative populations (Fig. 4.c). The hydrogen bonds are the same as proposed in ref. 29.

The most frequent hydrogen bond is E-Asp189, which is in place for more than 50 % of the snapshots for the complexes of fully- and unfluorinated variants Lys15TfeGly-BPTI and Lys15Abu-BPTI. For the partially fluorinated complexes of Lys15DfeGly-BPTI and Lys15MfeGly-BPTI, it occurs just higher than 30 %. Asp189 also accepts a hydrogen bond from B. This hydrogen bond is more frequent in the higher fluorinated complexes of Lys15TfeGly-BPTI and Lys15DfeGly-BPTI, where it is in place in 42 % of the snapshots, while it occurs for Lys15MfeGly-BPTI and Lys15Abu-BPTI 30 % and 35 %, respectively. The hydrogen bond D–C is again more often observed for the fully- and unfluorinated complexes. For Lys15MfeGly-BPTI, this can be explained by the fact of positions D and E being less frequently occupied. The other hydrogen bonds observed, namely A–B, B–A, B–C, C–B, B–E and C–E, are about equally frequent. Overall we do not observe a change in the structure of the water molecules in the S1 binding pocket or a trend in the stability of the hydrogen bond network with increasing fluorination.

To our surprise, we did not find any significant frequency of hydrogen bonds between the fluorine substituents and the water molecules at the hydration sites A-E. (We used the definition of a F-hydrogen bond from Ref. 48). The only exception is Lys15MfeGly-BPTI, in which the fluorine substituent occasionally forms hydrogen-bonds to a trapped water molecule (12.7 % of the frames), however in these cases the bonded water molecule cannot be assigned to any of the five hydration sites A-E. The absence of F-hydrogen bonds becomes plausible if one considers the orientations of the water molecules in the S1 binding pocket (Fig. 4.a). Water E acts as an hydrogen bond donor to Asp189 in *β*-trypsin and therefore faces with its oxygen towards BPTI. This enforces a similar orientation in the hydrogen-bonded water molecules A-D. Thus, the fluorine substituent, which would act as hydrogen bond acceptor, is in the vicinity of several water molecules, but the hydrogens in these water molecules face in the wrong direction.

#### c. Dynamics of the water molecules in the S1 binding pocket

Because the fluctuation around this static picture is rather large, we analyzed the dynamics of the water molecules in the binding pocket more closely. We calculated how long water molecules reside inside the S1 pocket, defined here as 5^Å^-spheres around the side chain ends of two critical amino acids at the bottom of the pocket, Asp189 and Ser190. The calculated survival probability is a measure of how likely it is, that a water molecule found in the S1 pocket at time *t_0_* will still be there at time *t_0_* +*t* (Fig. 5.a). After an initial fast decay until *t* ≈ 50 – 100ps, which reflects local fluctuations across the boundary of the 5Å-spheres, the survival probability decays exponentially with mean lifetimes of 0.9 to 1.3 ns. Water molecules in the Lys15Abu-BPTI-*β*-trypsin complex seem to have a longer residence time than in the fluorinated BPTI variants. Note, however, that the difference does not exceed one standard deviation and might not be statistically significant.

After determining the residence times for water molecules in the whole binding pocket, we were also interested in their mean residence time at the hydration sites A-E. As the water molecules which reside at these hydration sites exchange frequently throughout extensive MD simulations, keeping track of which water molecule resides at a specific hydration site for a given trajectory snapshot becomes a non-trivial task. We wrote a program to assign one water molecule for every simulation snapshot that resides at each of the hydration sites (see also section IV).

Fig. 5.b shows the average residence times of water molecules at the five positions A-E. In all four variants, water molecules at position A have a significantly higher average residence time than water molecules at any of the four other positions, likely due to its position deeply buried in the S1 pocket. Water molecules at positions B and D have low residence times of only a few 100 ps, and are thus highly dynamic. It is unlikely that they play a structurally important role. Specifically, water molecules B and D, which are not present in the wild-type BPTI-*β*-trypsin complex but are hypothesised to play a structural role in the fluorinated variant of the complex, have low residence times of 50 to 250 ps.

When comparing the fluorinated variants of BPTI, we find that residence times tend to increase with increasing fluorination for waters A-D, but not for water E. However, the effect is small compared to the standard deviation of the residence times. More importantly, the residence times in the fully fluorinated Lys15TfeGly-BPTI-*β*-trypsin are about as large as the residence times in the Lys15Abu-BPTI-*β*-trypsin complex. The partially fluorinated complexes show lower residence times.

In X-ray scattering experiments to determine the crystal structure of a protein, the thermal fluctuation reduces the intensity of the scattered X-rays. This effect is quantified it the so-called B-factors or Debye–Waller factor. Large B-factor imply large thermal motion of the corresponding atom in the crystal structure. The B-factors of the water molecules in the binding pocket of the fluorinated variants are considerably smaller than the water B-factors in the wild-type BPTI-*β*-trypsin complex (Fig. 3). In Ref. 29, it was proposed that this reduction in mobility is caused by stable hydrogen-bond network in the binding pocket of the fluorinated variants. The B-factors cannot easily be calculated from MD simulations. The closest proxy is the root-mean-square fluctuations (RMSF). When simulating the complex in water the water molecules are highly dynamic, frequently leave their position within the binding pocket and diffuse in and out of the binding pocket (See Fig. 5.a and 5.b). By contrast, the water molecules in the crystal structure are positioned at specific locations in the binding pocket. To nonetheless attempt a comparison between MD simulations and X-ray B-factors, we calculated the root-mean-square fluctuations (RMSF) of water molecules with a residence time longer than 200 ps at their respective hydration site. Fig. 3 shows the comparison. The water molecules in the five positions have similar RMSF values, where water molecules in position B have slightly lower RMSF values than the others. However, we do not find any trend in the RMSF with increasing fluorination. A possible explanation for this discrepancy might be that the reduced mobility of the water molecules in the crystal structures of the flourinated variants is caused by the larger volume of fluorine compared to hydrogen. In a crystal structure where the side-chains are fixed in one conformation, this reduces the space in which the water molecules can move. By contrast, in solution the binding pocket is not rigid and in particular the side chain of residue 15 rotates. This in turn would give the water molecules space to move explaining the similar RMSF values in all four variants of the BPTI-*β*-trypsin complex.

Overall, fluorination does not lead to a rigidification or stabilization of the water network in the S1 binding pocket. Instead partial fluorination (Lys15MfeGly and Lys15DfeGly), leads to an increased mobility of the water molecules, at least locally within the binding pocket. As in the structural analysis, we find that Lys15Abu and the fully fluorinated Lys15TfeGly show similar behaviour.

#### d. Energy and entropy of the water molecules in the S1 binding pocket

To evaluate the energy and entropy of the trapped water molecules we used Grid Inhomogeneous Solvation Theory (GIST)^55^ as implemented in the SSTMap^52–54^ command-line tool. In a GIST analysis, the two proteins in the complex are kept rigid while the water molecules are allowed to move freely. The simulation box is then discretized using a regular grid, where the grid cells are called voxels. The water-water (*E_ww_*) and the solute-water (*E_sw_*) interaction energy as well as the translational (*dS_tr_T*) and rotational (*dS_tr_T*) entropic part of the free energy at *T* = 300 K is calculated for each voxel. The entropic terms are calculated relative to the entropy in bulk water. Fig. 6 reports the Boltzmann-weighted averages of these four properties for each of the positions and for each of the four complexes. Neither the enthalpic nor the entropic terms show a trend with increasing fluorination. In particular the water-solute interactions of water molecules B and C which spacially replace the amino group of the lysine side chain do not change if the fluorination of the P1 methyl group changes. This indicates that the fluorinated methyl groups and the unfluorinated methyl group in Abu engage in similar interactions with these two water molecules and no specific interaction arises due to the fluorine substituents.

### E. Protonation state of His57

*β*-trypsin contains a histidine, His57, in the binding pocket which can be protonated at the two nitrogen atoms, N(*δ*) and N(*ϵ*), in its side chain. In the neutral form or unprotonated state, only one of the side chain nitrogen atoms is protonated, while in the protonated state both nitrogens are protonated. By itself, the pK_*a*_ of a protonated histidine side chain is 6.0^73^, and histidine is thus largely unprotonated at pH 7.

However, the pK_*a*_ of histidine residues can be varied drastically by the protein environment^74–76^. In the specific case of *β*-trypsin, His57 donates a hydrogen bond from N(*δ*) to the carboxyl group of Asp102. Since the N(*δ*)-proton is shared between His57 and Asp102, the equilibrium is shifted to the N(*ϵ*)-protonated state of His57 in the actual protein. That is the pKa of His57 is increased, and we expect it to be protonated. Therefore, the computational results presented so far were generated with His57 in the protonated state.

On the other hand the catalytic mechanism relies on a swift protonation and deprotonation of the His57 residue. We therefore additionally conducted simulation with His57 in the unprotonated state and tested the influence of the protonation state on the P1 residue and the water network.

Fig. 7 shows the *χ*_1_ torsion angle profiles of the P1 residue in all four Lys15X-BPTI-*β*-trypsin complexes. The unfluorinated Lys15Abu-BPTI and the fully fluorinated Lys15TfeGly-BPTI are not influenced by the protonation state of His57. The methyl group in Abu and the trifluoro-methyl group in TfeGly do not have an electric dipole moment and therefore do not interact with His57. Consequently, the P1 residues in Lys15Abu-BPTI and in Lys15TfeGly-BPTI have similar structural preferences. The *χ*_1_-torsion angle remains largely in the gauche+ conformation, i.e the conformation that is also found in all four crystal structures. The methyl group (Lys15Abu-BPTI) and the trifluoro-methyl group (Lys15TfeGly-BPTI) can rotate freely (rotation around *χ*_2_, data not shown) with equally populated gauche+, gauche- and trans conformations. The similar structure of the P1 residue in Lys15Abu-BPTI and Lys15TfeGly-BPTI might explain why the water network in these two variants tends to have similar structure and dynamics (Fig. 3 – Fig. 6).

The partially fluorinated variants, Lys15MfeGyl-BPTI and Lys15DfeGly-BPTI, can deviate from gauche+ conformation of *χ*_1_ torsion angle, which is found in the crystal structures. These side chains can possibly form dipole-dipole interactions with the backbone of a nearby loop or with the side chain of Ser195, which likely stabilizes the gauche- and the trans conformations. The conformational equilibrium between gauche+, gauche- and trans conformation is slightly influenced by the protonation state if His57 (Fig. 7).

We have also analyzed the water network in the binding pocket when His57 is unprotonated at N(*δ*). SI Fig. 4-8 in the supplementary material show the same analysis as presented in Figs. 3, 4.b, 4.c, 5.a, and 5.b for the protonated state. As in the protonated state, we do not find any trend that can explain the observed differences in the binding affinity. We do not find the water network and dynamics to be significantly influenced by the protonation state of His57.

Another concern is whether Asp189 at the bottom of the binding pocket might be protonated. For uncomplexed trypsin, Czodrowski et al.^77^ calculated the pKa of Asp189 using their own modification of the PEOE method^78^ and found it to be 4.3, whereas a PROPKA^79,80^ calculation yielded 6.29^81^. However, these differences do not necessarily show differences in the methods, but might also be caused by the different conformations that were used for these estimates. In previous work (Fig. 11 in Joswig et al. 20 2 1 ^76^), we calculated the pKa distribution of aspartates and glutamates for a carbohydrate recognition receptor using PROPKA and found that the pKa of a single carboxyl group can vary by four pKa units - depending on the conformation. Overall, we do not believe that Asp189 is protonated to a significant extent in the ensemble, because *β*-trypsin recognizes positively charged side-chains using the negative charge on the unprotonated Asp189^9^. Thus, *β*-trypsin wouldn’t be target specific if Asp189 was protonated. Additionally, Asp189 is partially solvated in the binding pocket, and solvent exposed aspartates are unlikely to be protonated^82^ and usually need a second carboxyl group in close vicinity to stabilize the protonated state, like in Refs. 76, 83, and 84. *β*-trypsin does not have a second carboxyl group in the binding pocket.

### F. Helical propensity of the P1 residue

Fluorinating the P1 residue could influence the local structure around the P1 residue of the complex and thereby influence the stability of the complex. Fig. S9 in the supplementary material shows the Ramachandran plots of Abu, MfeGyl, DfeGly, and TfeGly in a the dipeptide (i.e. Ace-X-Nme with X = Abu, MfeGyl, DfeGly, TfeGly), in the P1 position of uncomplexed BPTI, and in the P1 position of BPTI in complex with *β*-trypsin. In the dipeptides, we observe a slight decrease in the population of the *α*-helical state with increasing fluorination. In uncomplexed BPTI we observe the opposite trend. Here we find a large population in the *α*-helical state in all four variants, and slightly shifted *β*-sheet state which is stabilized with increasing fluorination, this depleting the *α*-helical state. In the complexes, the *α*-helical state and the *Lα*-helical state is populated, and three of the four complexes additionally show some population in the *β*-sheet state. However, no clear trend for the influence of the fluorination can be observed. Overall, we find that fluorination has some influence on the local structure, but the effects seem to be small and with the current data it is not obvious how these local changes influence the stability of the complexes.

## IV. CONCLUSIONS

We provide insights from experiments and computational simulation into the water network in the trypsin S1 binding pocket, in complex with Lys15Abu-BPTI and fluorinated analoga. We repeated the inhibition assay from Ref. 29, including the now synthetically available Lys15MfeGly-BPTI, and found a step-wise increase of inhibitor strength with increasing fluorination. In MD simulations, using umbrella sampling and our self-parametrized force field for fluorinated amino acids, we confirmed this step-wise increase in inhibitor strength and estimated a similar binding free energy. We determined the X-ray crystal structure of the complex of trypsin with Lys15MfeGly and found the pocket water configuration to be similar as in the complexes of Abu and the di- and tri-flourinated analoga.

Following the hypothesis of fluorine forming interactions with water molecules and thereby rigidifying the water network in the S1 binding pocket, we analyzed the structure and dynamics of the S1 water molecules computationally with MD simulations. We found the positional water structure seen in the crystal structure to be on average consistent with the protein crystal structure, but we observe the water molecules frequently exchanging positions and leaving or entering the binding pocket.

The water molecules form a hydrogen bond network that is similar for trypsin in complex with Lys15Abu-BPTI and its fluorinated analoga. Surpisingly, we do not find hydrogen bond like interactions between the fluorinated methyl groups and water molecules at the hydration sites A-E. We estimate a water molecule to stay in the S1 binding pocket for a mean lifetime of 0.9-1.3 ns and the exchanges at the specific hydration sites to be even faster, with mean residence times of 50-250 ps with no trend in regard to fluorination. Moreover the enthalpic and entropic properties of water in the region of the hydration sites also do not indicate increasing stability of water molecules with increasing fluorination.

We additionally tested whether the protonation state of His57 in the binding pocket has an influence on the water network. Just as in the protonated state, we did not find any trend when comparing the unprotonated Lys15Abu-BPTI to the partially fluorinated variants Lys15MfeGly-BPTI and Lys15DfeGly-BPTI and the fully fluorinated Lys15TfeGly-BPTI.

We conclude that we could not observe a rigidification or stabilizing effect of fluorination on the water network in the S1 pocket. Ref. 71 reports the van-der-Waals volumes of Abu and its fluorinated variants, which monotonously increase from *V*_VdW_ ≈ 40 Å^3^ in Abu to *V*_vdW_ ≈ 60 Å^3^ in TfeGly. The corresponding decrease of the B-factors of the water molecules in the X-ray crystal structure (Fig. 3.b) is thus possibly caused by a steric effect and not by the stabilization of the hydrogen bond network. Overall, the experimentally and computationally observed increase in inhibitor strength, respectively binding free energy, cannot be explained by a change in the structure or the dynamics of the water network. Other factors have to play a role.

Ref. 33 investigated the hydrophobicity of fluorinated residues and reports an increase of the hydration free energy from Abu to the fully fluorinated Lys15TfeGly of 1 kcal/mol = 4.2 kJ/mol. Since the P1 residue in BPTI is solvent-exposed, this effect will certainly affect the hydration free energy of the fluorinated BPTI-variants. Thus, the observed increase in inhibitor strength might partially be due to an increase in the free energy of the unbound BPTI, rather than a decrease in the free-energy of the bound state. But we do not think it can exclusively be attributed to the difference in hydrophobicity, since the P1 side chain is (partially) hydrated even in the binding pocket.

Furthermore, while the transition region between bound and unbound state is likely not fully converged in our PMFs, we observe an encounter complex (ξ > 3.0nm) in all four systems. The stability of this encounter complex seems to increase with increasing fluorination. A similar encounter complex has recently been discovered in the binding process of wt-BPTI to *β*-trypsin^19^. Additionally, we observe a pre-bound intermediate state whose stability sensitively depends on the fluorination. This pre-bound state does not seem to exist in the wt-BPTI-*β*-trypsin complex^19^. We plan to investigate the free energy landscape and the unbinding process more closely using dynamic reweighting techniques^85–87^.

From a broader perspective, our study shows that fluorination is a sensitive tool to tune the affinity of a protein-protein complex. However, one cannot rationalize the observed effects by only considering the fully bound complex. Rather fluorine substituents also influence the solvation properties of the unbound ligand, the specific interactions within the fully bound complex, the stability of the encounter complex and the stability of intermediate states along the binding pathway. All these effects act together to generate the affinity of a fluorinated protein-protein complex.

## Supporting information

Force Field Files

Crystallographic Data

PDB File

V.

## ACKNOWLEDGEMENTS

Gefordert durch die Deutsche Forschungsgemeinschaft (DFG): Projektnummer 387284271 – SFB 1349, Projektnummer 434130070 - GRK 2662. Funded by the Deutsche Forschungsgemeinschaft (DFG, German Research Foundation):Project-ID 387284271 – SFB 1349, Project-ID 434130070 - IRTG 2662. We acknowledge access to high-performance computers via the Zentraleinrichtung fur Datenverarbeitung (ZEDAT) of Freie Universität Berlin. We acknowledge access to beamlines of the BESSY II storage ring (Berlin, Germany) via the Joint Berlin MX-Laboratory sponsored by the Helmholtz Zentrum Berlin fuär Materi-alien und Energie, the Freie Universitaät Berlin, the Humboldt-Universitäat zu Berlin, the Max-Delbriick-Centrum, the Leibniz-Institut fur Molekulare Pharmakologie and Charité – Universitäatsmedizin Berlin.

## VI. SUPPLEMENTARY MATERIAL

- Amended force field AMBER14sb (amber14sb_amended.ff.zip) and plots of the partial charges and Lennard-Jones parameters (SI.pdf)
- Structure and dynamics of water in the *β*-trypsin S1 binding pocket for unprotonated His57 (SI.pdf)
- Crystallographic data (crystallographic_data.docx and SI.pdb)
- Code repository: https://github.com/bkellerlab/tracffiwaters

## Notes

https://github.com/bkellerlab/track_waters

### Competing Interest Statement

The authors have declared no competing interest.

### Summary of Updates

Revision according to peer-review process has been updated.

