## Supplementary Information for "Water Network in the Binding Pocket of Fluorinated BPTI-Trypsin Complexes - Insights from Simulation and Experiment"

### I. PARAMETRIZATION OF ABU AND FLUORINATED DERIVATIVES.

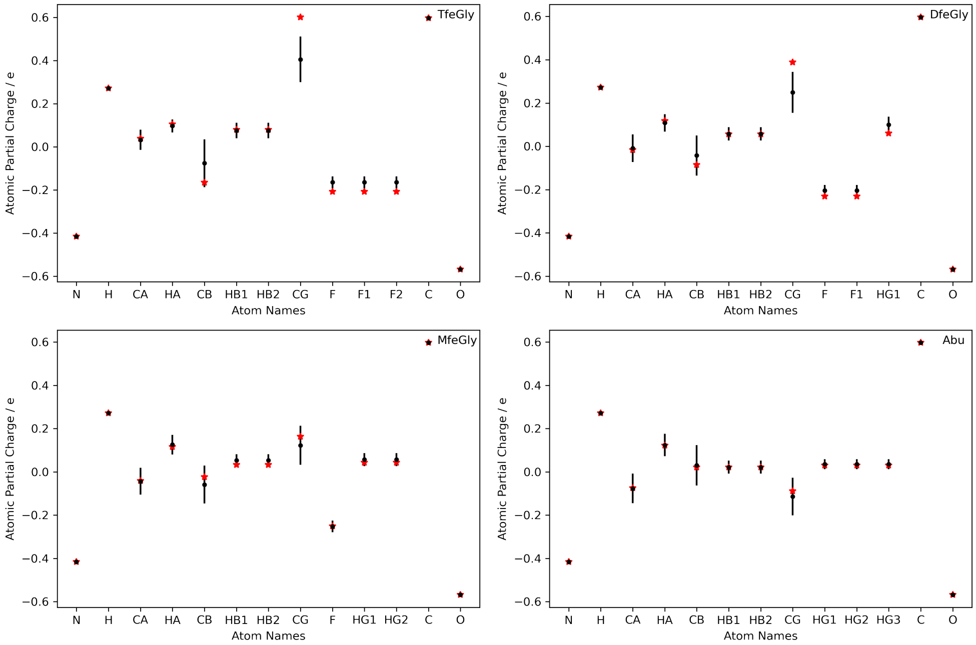


Figure S1. Charges of TfeGly, DfeGly, MfeGly and Abu in our forcefield (black), compared to those of Robalo et al.[1,2] (red). The error bars indicate standard deviation of the charges after the multiconformational fit.

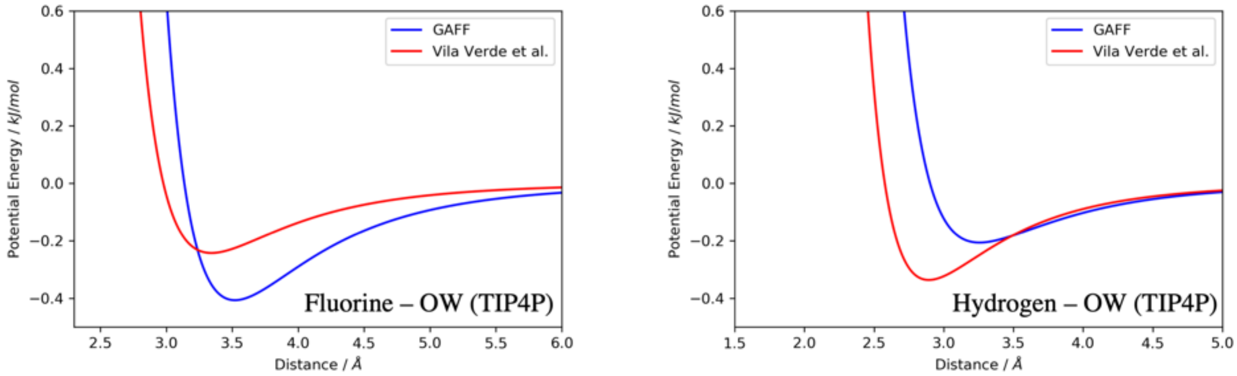


Figure S2. Lennard-Jones interaction potential of Fluorine and hydrogen of the fluorinated methyl group $H_F$. The blue profile is calculated from the General Amber Force Field (GAFF) parameters, the red profile is calculated from the parameters of Ref. 1 and 2.

#### II. BOOTSTRAPPING OF UMBRELLA SAMPLING SIMULATIONS.

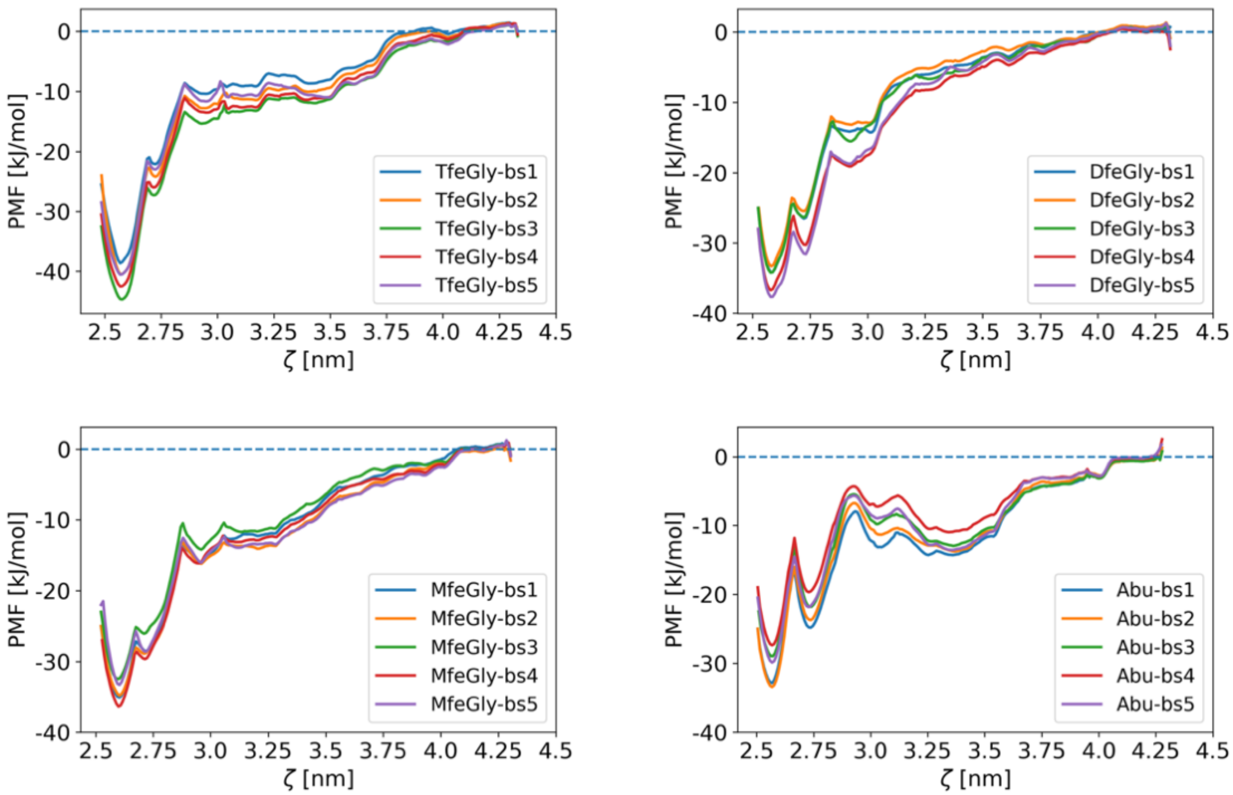


Figure S3. PMF profiles of samples from simplified bootstrapping of umbrella sampling simulations. The 50 ns windows were separated into 10 ns trajectories. Bootstrapping samples are following combinations: bs1 (0-10, 10-20, 20-30, 30-40); bs2 (0-10, 10-20, 20-30, 40-50); bs3 (0-10, 10-20, 30-40, 40-50); bs4 (0-10, 20-30, 30-40, 40-50); bs5 (10-20, 20-30, 30-40, 40-50).

#### III. UNPROTONATED STATE OF HIS57

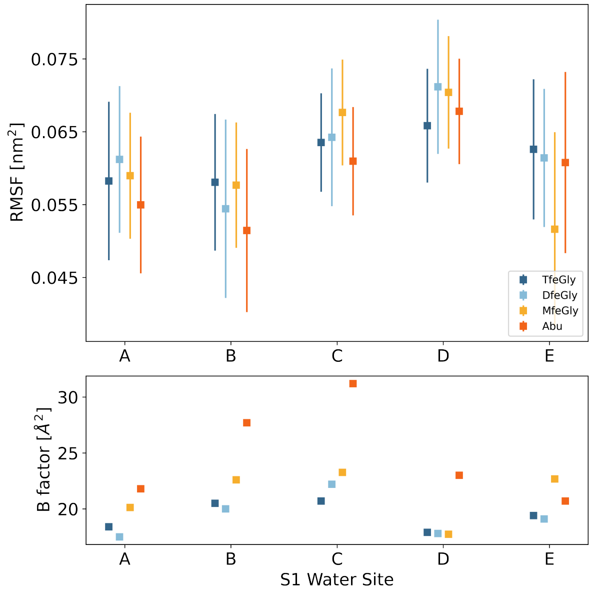


Figure S4. Figure corresponding to Fig. 3 in the main text but for the unprotonated state of His57. (top) RMSF of water molecules with a lifetime longer than 200 ps at the hydration sites A-E in the $\beta$-trypsin S1 binding pocket. (bottom) B factors of the waters in the $\beta$-trypsin S1 binding pocket as reported in Ref. 3.

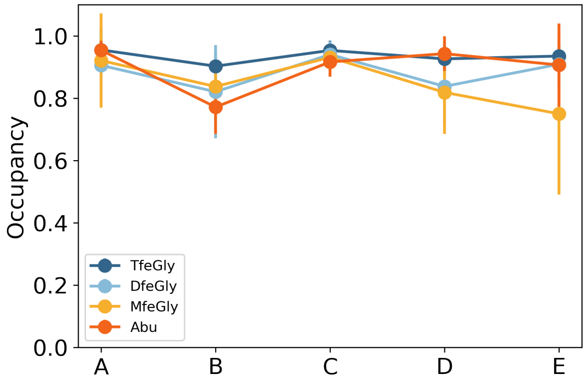


Figure S5. Figure corresponding to Fig. 4b in the main text but for the unprotonated state of His57. Structure of the water molecules in the S1 binding pocket: relative populations of the hydration sites A-E.

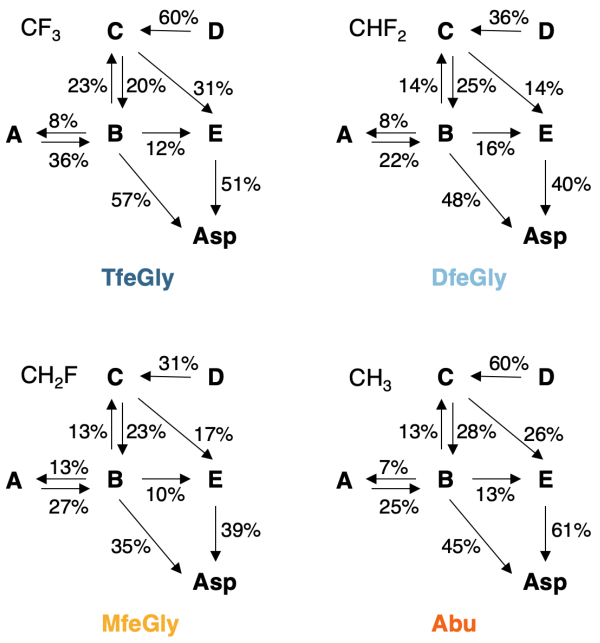


Figure S6. Figure corresponding to Fig. 4c in the main text but for the unprotonated state of His57. Structure of the water molecules in the S1 binding pocket: hydrogen bond network. Relative populations of the hydrogen bonds formed by waters A-E to each other and to the side chain of Asp189. The arrows point from the donor to the acceptor.

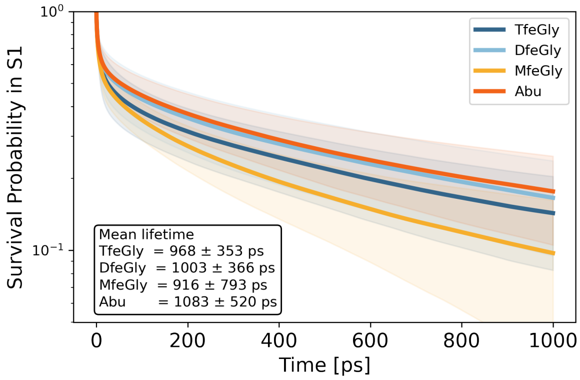


Figure S7. Figure corresponding to Fig. 5a in the main text but for the unprotonated state of His57. Dynamics of the water molecules in the S1 binding pocket: average survival probability a water molecule in the binding pocket. The mean lifetime is approximated by fitting a single-exponential decay function to the data after the initial burn phase.

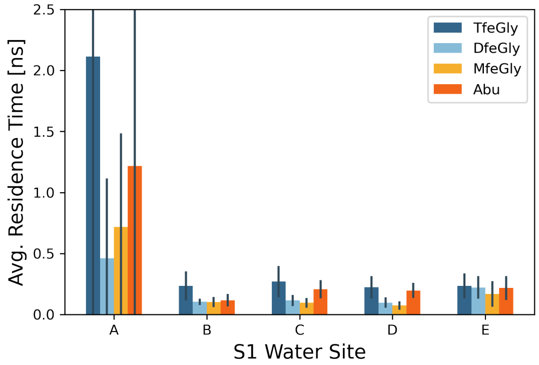


Figure S8. Figure corresponding to Fig. 5b in the main text but for the unprotonated state of His57. Dynamics of the water molecules in the S1 binding pocket: residence time of a water molecule at the sites A-E. Data analyzed form 1 µs of simulation data for each system.

##### A. Computational Details

The simulations with the unprotonated state of His57 of $\beta$-trypsin were set-up, equilibrated and run in the same way as the unbiased simulations with the protonated state of His57 described in the method section of the main text. When setting up the starting configuration and topology with Gromacs, N($\delta$) of was chosen to be unprotonated, while N($\epsilon$) was kept protonated in what we here call the unprotonated state.

#### IV. BACKBONE CONFORMATION OF ABU AND FLUORINATED DERIVATIVES

The backbone conformations of Abu and its fluorinated derivatives MfeGly, DfeGly and TfeGly were compared as a dipeptide (Ace-[Abu-variant]-Nme), in free BPTI and in the BPTI-$\beta$-trypsin complex. MD simulations were run and the $\phi$ and $\psi$ backbone dihedrals were calculated throughout the simulation. The simulations were set-up and equilibrated in the same way as described for the unbiased simulations in the main text. For the dipeptide, 1 µs of simulation time per derivative was calculated, for the free BPTI and complexes 100 ns each was calculated.

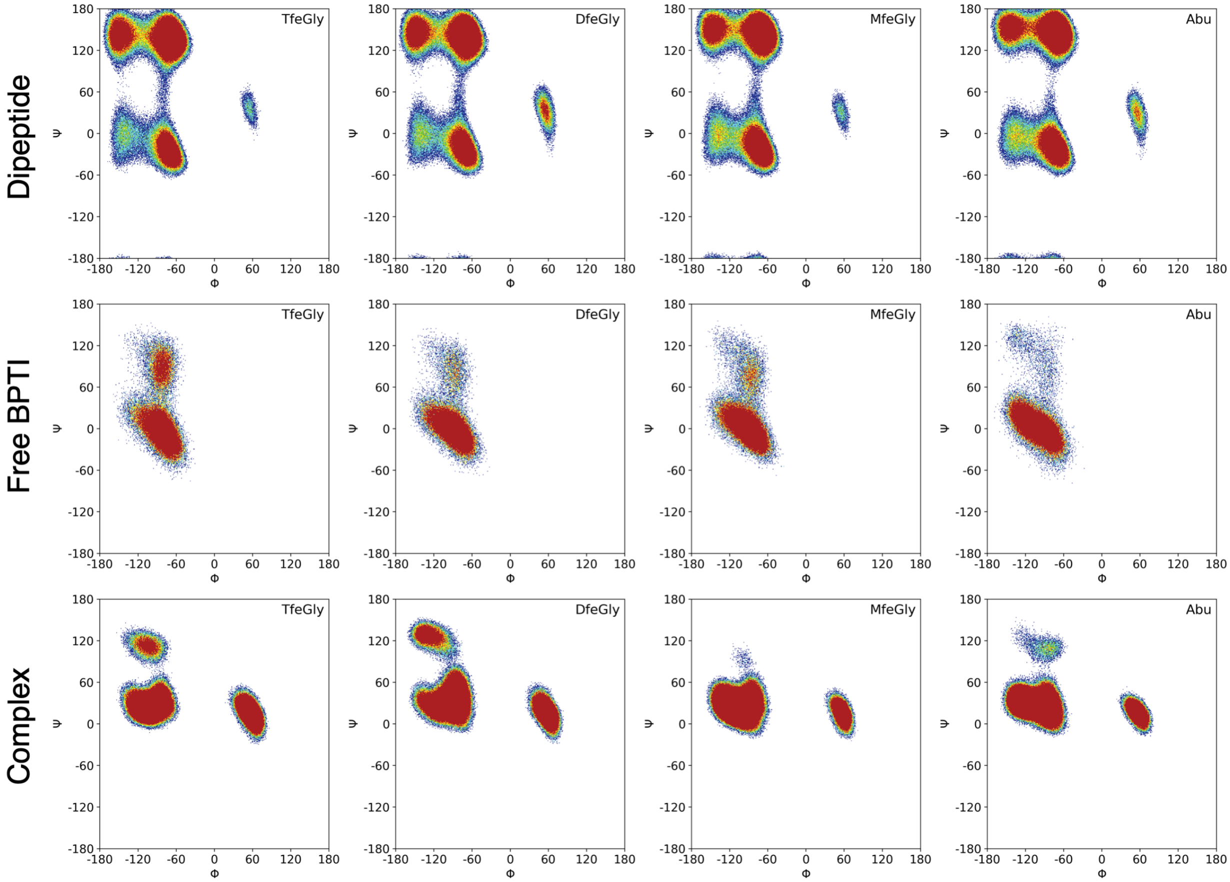


Figure S9. Ramachandran plots of Abu derivatives as dipeptide (Ace-[Abu-variant]-Nme; top row), in free BPTI (middle row) and in BPTI in complex with $\beta$-trypsin (bottom row) from MD simulations.

#### V. CONVERGENCE OF THE UNBOUND STATE IN THE PMF CALCULATIONS

##### A. Sampling of Diffusion of the Unbound BPTI Derivative in the Umbrella Windows

We measured how much of the space around $\beta$-trypsin is explored by the BPTI derivative in the respective umbrella windows of the unbound state. The trajectores of the umbrella windows were superposed on the backbone of $\beta$-trypsin and the COM of $\beta$-trypsin was moved to the origin of the coordinates for each simulation snapshot. The position of the COM of the BPTI derivative was then transformed into spherical coordinates. This transformation allowed a projection of the BPTI derivative positions on the surface of a sphere, because the radial coordinate, i.e. the COM distance between the proteins, is constrained by the umbrella potential. The positions of the COM of the BPTI derivative (from the azimuthal and polar coordinate) were then plotted on the surface of a sphere. The hammer projection of such a surface plot is shown below in Fig. S10 for one exemplary umbrella window (at $\xi \approx 4.25\,\text{nm}$) in the unbound state. For comparison, the same analysis for an umbrella window in the fully bound state is shown in Fig. S11.

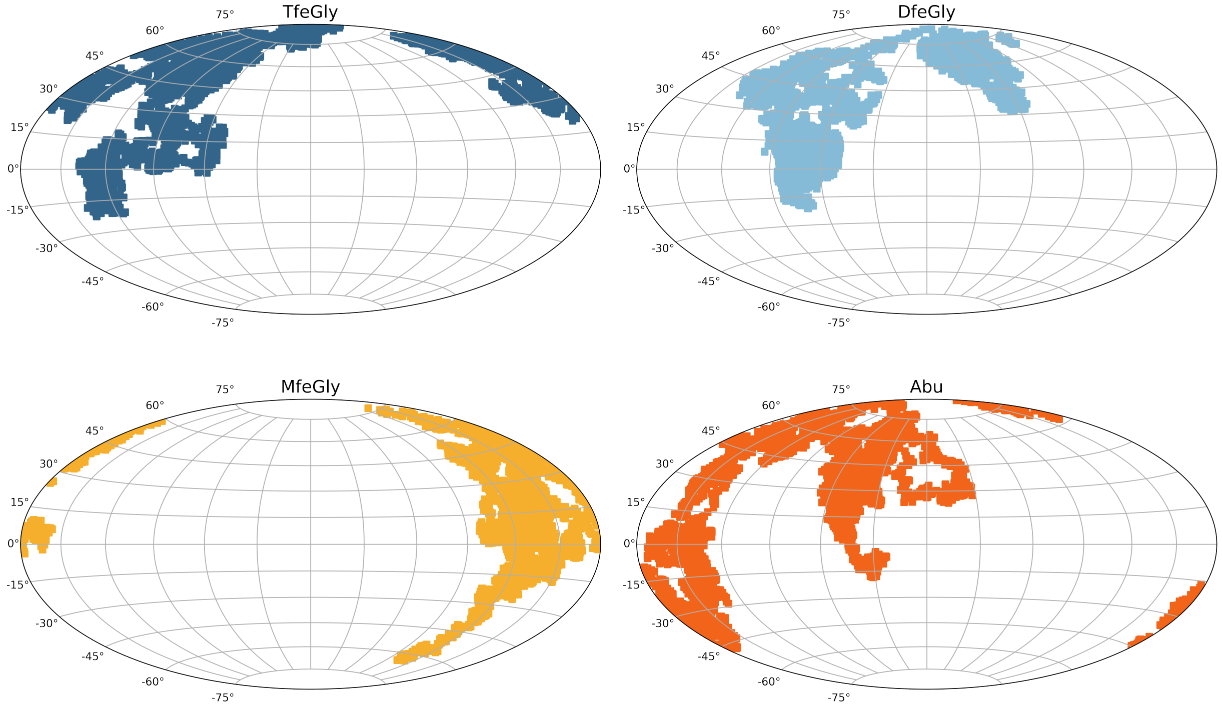


Figure S10. Positions of the COM of the BPTI derivative that are sampled in the unbound state, exemplary in umbrella 34 at $\xi \approx 4.25\,\text{nm}$, relative to the COM of $\beta$-trypsin.

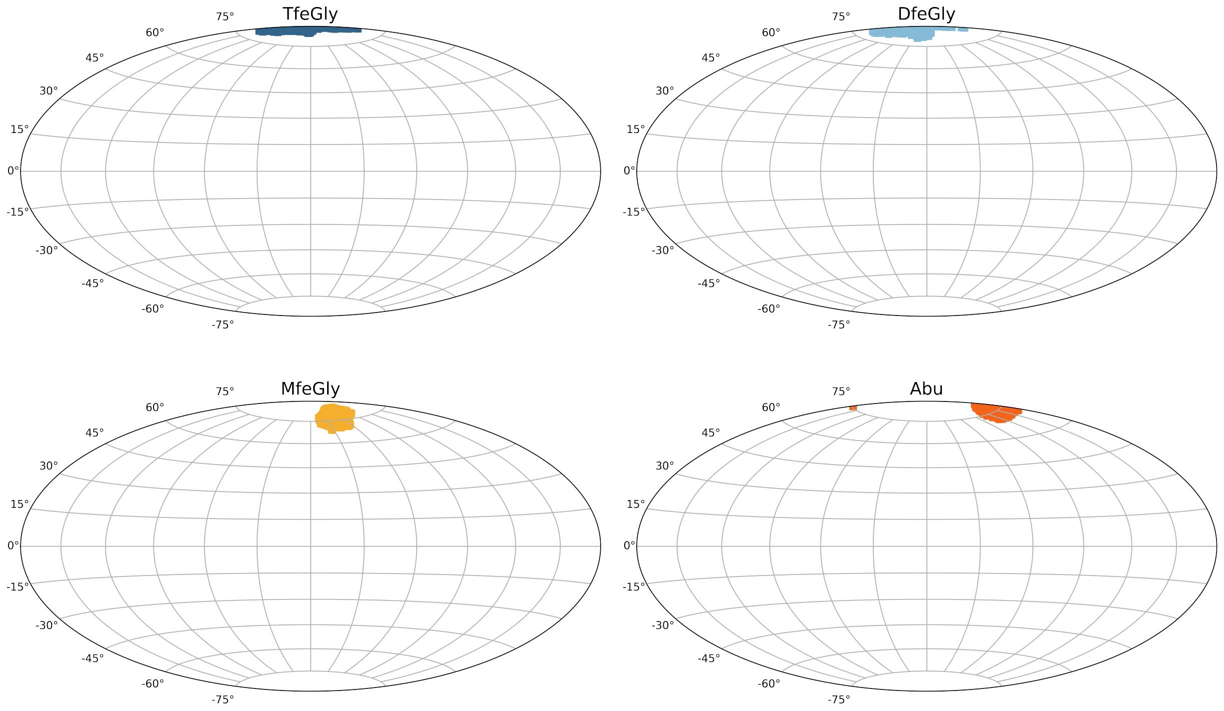


Figure S11. Positions of the COM of the BPTI derivative that are sampled in umbrella 1, representative for the fully bound complex relative to the COM of $\beta$-trypsin.

#### B. Relative orientation Between Trypsin and BPTI Derivatives in the Bound and Unbound State

The relative orientation between $\beta$-trypsin and the BPTI derivatives was measured in the respective umbrella windows of the fully bound complex and the unbound state. The orientation angle is calculated by defining a plane on the surface of $\beta$-trypsin and a vector along the long axis of the BPTI derivative. Then, the vector orthogonal to the plane is defined and the orientation angle was then calculated as the angle between this orthogonal vector and the vector along BPTI. This calculation was conducted for every simulation snapshot in the respective umbrella windows and the distribution is shown in violin plots below in Fig. S12 for the fully bound complex and in Fig. S13 for the unbound state. The plane on the surface of $\beta$-trypsin is defined by the $C_\alpha$ atoms of Asp196, Gly198 and Trp217. These amino acids were chosen so the plane lies approximately flat on the S1 pocket. The vector along the BPTI derivative was defined by the $C_\alpha$ atoms of the P1 residue and Phe22.

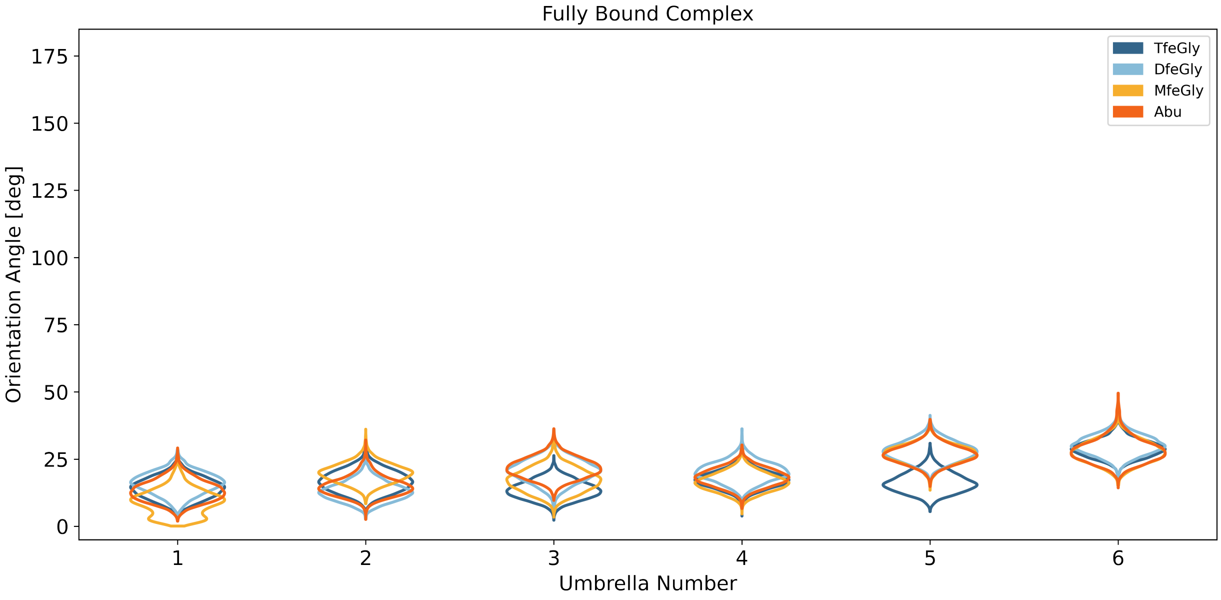


Figure S12. Relative orientation of the BPTI derivatives to $\beta$-trypsin in the bound state ($\xi < 2.8\,\mathrm{nm}$).

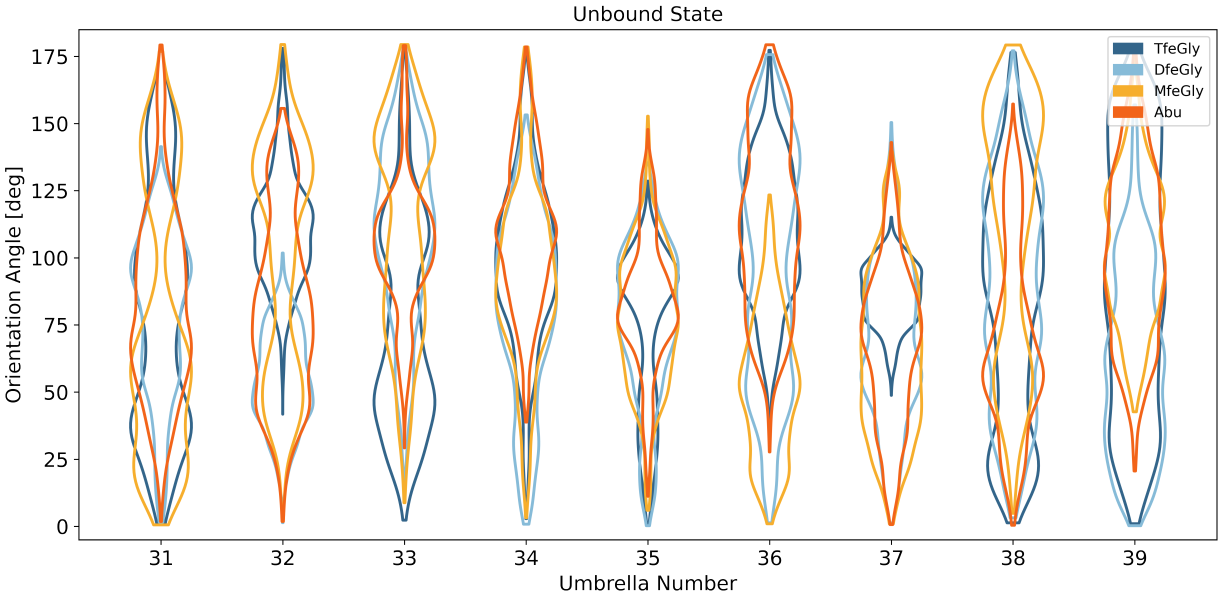


Figure S13. Relative orientation of the BPTI derivatives to $\beta$-trypsin in the unbound state ($\xi > 4.1\,\mathrm{nm}$).

##### C. Discussion

In order to calculate absolute free-energy differences from PMFs the rotational and translational sampling in the unbound state needs to be taken into account accurately. The unbound state is fully sampled if

1. $\beta$-trypsin and the BPTI variant are sufficiently far apart that the BPTI variant can diffuse freely.

2. The tumbling of BPTI (i.e. the rotation around its axes of inertia) is fully sampled.

3. The diffusion the spherical volume at at given value of the reaction coordinate is fully sampled.

At a distance of $\xi > 4.1$ nm, we can be confident that condition 1 is fulfilled. Also Figs. S13 and S10 indicated that this is the case. Fig. S13 shows that from umbrella 31 on, the BPTI variants can tumble freely and in most umbrellas all orientations are sampled (exception are umbrella 35 and 37). Thus, we argue that condition 2 is sufficiently fulfilled. Fig. S10 shows that the spherical volume is not fully sampled and condition 3 is violated. Given a simulation time of 50 ns per umbrella this is expected. However, the four BPTI-variants are extremely similar in mass and shape. Thus, their diffusive properties should be almost identical. Within a given umbrella window they then sample approximately the same volume, and even with partial sampling, the unbound states of the four systems should be comparable. Fig. S10 seems to confirm this assumption: all four systems sample approximately $\frac{1}{3}$ of the spherical volume. The PMFs can thus serve as a rough test as to whether the MD simulations capture the experimental trends. For quantitative free energy differences, additional sampling is needed.
