## Supplementary material for "Water Network in the Binding Pocket of Fluorinated BPTI-Trypsin Complexes - Insights from Simulation and Experiment": Crystallographic Data

Table 6.4. Crystallographic data collection, refinement, and validation statistics.

| **Dataset** | Trypsin/BPTI MfeGly |
| --- | --- |
| PDB entry | 7PH1 |
| **Data** **Collection** | |
| Wavelength [Å] | 0.9184 |
| Temperature [K] | 100 |
| Space group | *I*222 |
| Unit Cell Parameters  a, b, c [Å]  α, β, γ [°] | 74.97, 81.29, 124.25  90.0, 90.0, 90.0 |
| Resolution [Å]^a^ | 30.00 - 1.18  (1.25 - 1.18) |
| Reflections ^a^  Unique ^a^  Completeness [%]^a^  Multiplicity ^a^ | 124,357 (19,736)  99.8 (98.9)  7.4 (7.5) |
| Data quality ^a^  Intensity [I/σ(I)] ^a^  R_meas_ [%]^a ,b^  CC_1/2_ ^a,c^  Wilson B value [Å^2^] | 10.02 (0.91)  9.3 (204.9)  99.9 (45.6)  19.0 |
| **Refinement** | |
| Resolution [Å]^a^ | 30.00 - 1.18  (1.21 - 1.18) |
| Reflections ^a^  Number  Test Set [%] | 124,220  1.5 |
| R_work_ [%]^a^  R_free_ [%]^a^ | 14.9 (34.3)  16.6 (35.8) |
| Asymmetric Unit  Protein: Residues, Atoms  Ligands: Molecules  Water molecules | 223 (E), 3,693 (E)  57 (I), 1,029 (I)  1 (Ca^2+^), 8 (glycerol),  14 (SO_4_^2-^)  322 |
| Mean Temperature factors [Å^2^]^b^  All Atoms  Macromolecules  Ligands  Water molecules | 22.9  23.8 (E), 20.5 (I)  23.2 (Ca^2+^), 36.9 (glycerol)  36.5 (SO_4_^2-^)  32.4 |
| RMSD from Target Geometry ^d^  Bond Lengths [Å]  Bond Angles [°] | 0.015  1.512 |
| **Validation Statistics** | |
| Ramachandran Plot ^f^  Residues in Allowed Regions [%]  Residues in Favored Regions [%]  Ramachandran plot *Z*-score ^f^ (RMSD)  whole  helix  sheet  loop  Molprobity Clashscore ^g^  Molprobity score ^f^ | 1.5  98.5  -0.25 (0.41)  1.92 (0.74)  -0.97 (0.49)  -0.20 (0.39)  1.62  0.88 |

^a^	data for the highest resolution shell in parenthesis

^b^	R_meas_(I) = ∑_h_ [N/(N-1)]^1/2^ ∑_i_ │I*_i_*_h_ - <I_h_>│ / ∑_h_∑_i_ I*_i_*_h_, in which <I_h_> is the mean intensity of symmetry-equivalent reflections h, I*_i_*_h_ is the intensity of a particular observation of h and N is the number of redundant observations of reflection h. ^2^

^c^	CC_1/2_ = (<I^2^> - <I>^2^) / (<I^2^> - <I>^2^) + σ^2^_ε_, in which σ^2^_ε_ is the mean error within a half-dataset.^3^

^d^	RMSD – root mean square deviation

^e^	calculated with PHENIX ^4^
^f^	calculated with MOLPROBITY ^1^
^g^	Clashscore is the number of serious steric overlaps (> 0.4 ) per 1,000 atoms.^7^

1.	C. J. Williams, J. J. Headd, N. W. Moriarty, M. G. Prisant, L. L. Videau, L. N. Deis, V. Verma, D. A. Keedy, B. J. Hintze, V. B. Chen, S. Jain, S. M. Lewis, W. B. Arendall, 3rd, J. Snoeyink, P. D. Adams, S. C. Lovell, J. S. Richardson and D. C. Richardson, *Protein Sci*, 2018, **27**, 293-315.

2.	K. Diederichs and P. A. Karplus, *Nat. Struct. Biol.*, 1997, **4**, 269-275.

3.	P. A. Karplus and K. Diederichs, *Science*, 2012, **336**, 1030-1033.

4.	P. D. Adams, P. V. Afonine, G. Bunkoczi, V. B. Chen, I. W. Davis, N. Echols, J. J. Headd, L. W. Hung, G. J. Kapral, R. W. Grosse-Kunstleve, A. J. McCoy, N. W. Moriarty, R. Oeffner, R. J. Read, D. C. Richardson, J. S. Richardson, T. C. Terwilliger and P. H. Zwart, *Acta crystallographica. Section D, Biological crystallography*, 2010, **66**, 213-221.
